# Impairment of the SKN-1A/NRF1 proteasome surveillance pathway triggers tissue-specific protective immune responses against distinct natural pathogens in *C. elegans*

**DOI:** 10.1101/2023.08.22.554255

**Authors:** Manish Grover, Spencer S. Gang, Emily R. Troemel, Michalis Barkoulas

## Abstract

Protein quality control pathways play important roles in resistance against pathogen infection. For example, the conserved transcription factor SKN-1/NRF upregulates proteostasis capacity after blockade of the proteasome, and also promotes resistance against bacterial infection in the nematode *C. elegans*. SKN-1/NRF has three isoforms, and the SKN-1A/NRF1 isoform in particular regulates proteasomal gene expression upon proteasome dysfunction as part of a conserved bounce-back response. We report here that, in contrast to the previously reported role of SKN-1 in promoting resistance against bacterial infection, loss-of-function mutants in *skn-1a* and its activating enzymes *ddi-1* and *png-1,* show constitutive expression of immune response programmes against natural eukaryotic pathogens of *C. elegans*. These programmes are the Oomycete Recognition Response (ORR), which promotes resistance against oomycetes that infect through the epidermis, and the Intracellular Pathogen Response (IPR), which promotes resistance against intestine-infecting microsporidia. Consequently, *skn-1a* mutants show increased resistance to both oomycete and microsporidia infections. We also report that almost all ORR/IPR genes induced in common between these programmes are regulated by the proteasome and interestingly, specific ORR/IPR genes can be induced in distinct tissues depending on the exact trigger. Furthermore, we show that increasing proteasome function significantly reduces oomycete-mediated induction of multiple ORR markers. Altogether, our findings demonstrate that proteasome regulation keeps innate immune responses in check in a tissue-specific manner, against natural eukaryotic pathogens of the *C. elegans* epidermis and intestine.

## INTRODUCTION

The evolutionary history of *Caenorhabditis elegans* is shaped by various biotic interactions in its natural habitat [1]. These interactions include beneficial and pathogenic microbes, which trigger a diverse set of responses in *C. elegans* [1]. Therefore, studying *C. elegans* under some conditions encountered in its natural environment can provide novel insights, such as new functions for genes lacking functional annotation in the genome [1]. For example, the identification of oomycetes as natural pathogens of *C. elegans*, revealed a previously uncharacterized family of *chitinase-like (chil)* genes as immune response effectors, which can modify cuticle properties to prevent oomycete attachment and consequently infection [2]. Most *chil* genes are not expressed in lab culture conditions and can only be induced upon pathogen recognition together with a broader subset of genes previously described as the oomycete recognition response (ORR) [3]. *C. elegans* can similarly mount specific responses against other natural pathogens as well, for example, the intracellular pathogen response (IPR) against microsporidia and the Orsay virus [4, 5].

While the nematode is likely to use its sensory capabilities to detect pathogens and activate pathogen-specific responses [6], it also uses surveillance immunity as a broad way to tackle pathogenic attacks [7]. Pathogens can hijack the host cellular machinery to support their growth and development, and in doing so they may disrupt core cellular processes, such as transcription, translation, protein turnover and mitochondrial respiration. As a result, disruption to these processes can be sensed as a pathogenic attack leading to activation of immune responses [8–10]. For example, inhibition of translation by *Pseudomonas aeruginosa* exotoxin A leads to activation of immune response genes through transcription factors such as ZIP-2 and CEBP-2 [8, 9, 11]. Mitochondrial disruption by *P. aeruginosa* also induces immune responses [12, 13]. Similarly, RNAi and microbial toxin perturbation of various core cellular processes induces detoxification enzymes and aversion behaviour in *C. elegans* [10]. Likewise, the IPR can be induced either by inhibition of the purine salvage pathway as seen in *pnp-1* mutants [14] or by blockade of the major protein degradation machinery in the cell, the proteasome [4, 15].

A major player regulating proteostasis upon proteasome blockade is the conserved transcription factor SKN-1/NRF [16]. One of its isoforms, SKN-1A (NRF1 in humans), is normally localized in the endoplasmic reticulum membrane where it is glycosylated and then targeted for proteasomal degradation (Figure 1A) [17]. However, upon proteasome blockade, the glycosylated isoform escapes degradation and is edited by PNG-1 (NGLY in humans), a conserved N-glycanase that converts the N-glycosylated asparagine residues to aspartic acid, and the edited protein then translocates to the nucleus [17]. Inside the nucleus, DDI-1 (DDI2 in humans), a conserved aspartic protease, cleaves the N-terminal region of the protein, and this processed SKN-1A is now activated to upregulate expression of proteasomal subunits to increase proteostasis capacity in a bounce-back response (Figure 1A) [17–19]. In addition to its role in promoting proteostasis capacity, SKN-1 is also required to respond to oxidative stress [20] and promotes resistance to bacterial pathogens, such as the human pathogens *P. aeruginosa* and *Enterococcus faecalis* [21–23].

**Figure 1:**
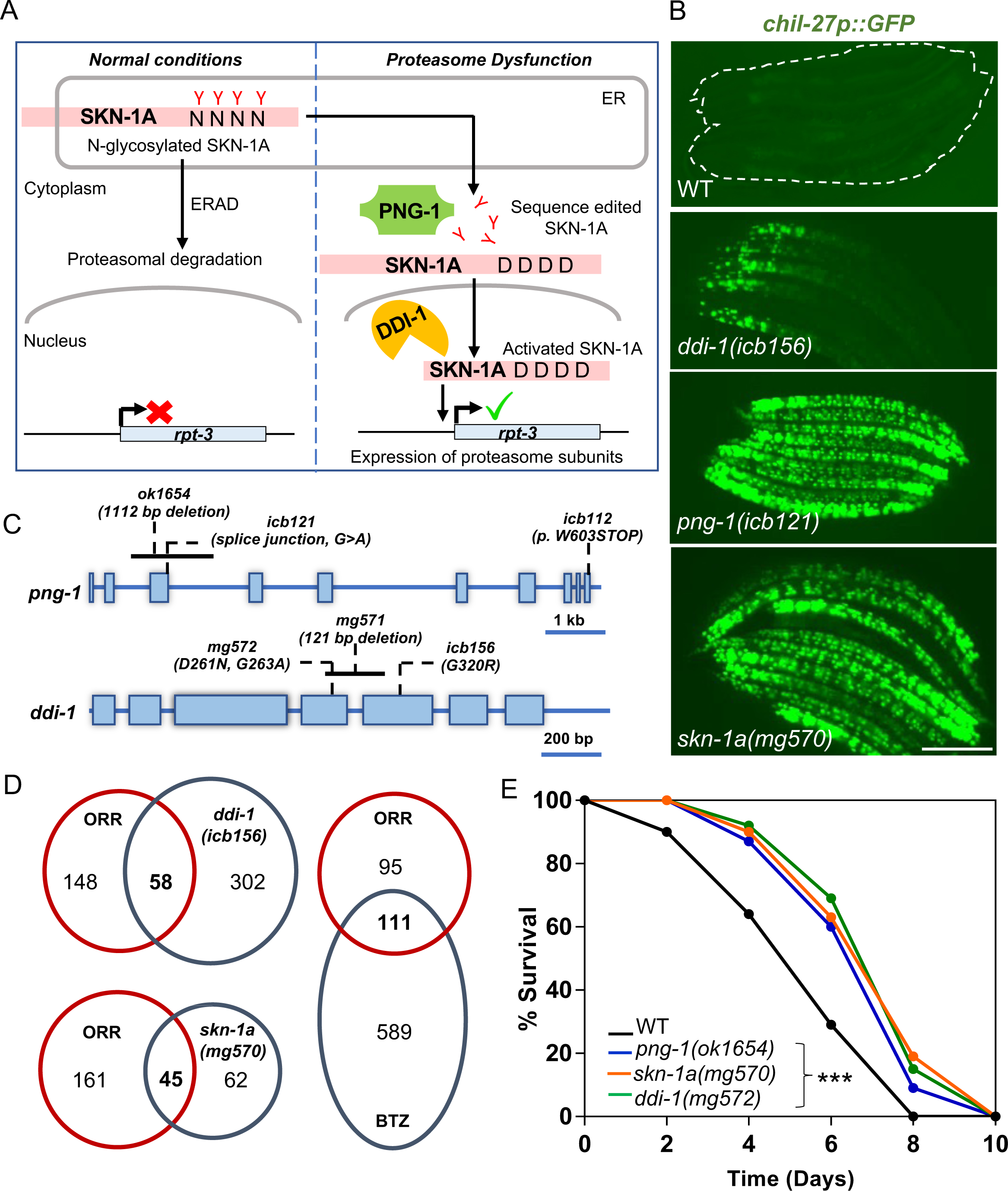
Proteasome impairment activates oomycete recognition response (ORR) in *C. elegans*. **(A)** Schematic showing the proteasome surveillance pathway and the stepwise activation of SKN-1A for transcription of proteasomal subunits. **(B)** L4 stage *C. elegans* showing constitutive *chil-27p::GFP* expression in *ddi-1(icb156)*, *png-1(icb121)*, and *skn-1a(mg570)* mutant animals. Note that *skn-1a* and *png-1* show a stronger, full body activation of the *chil-27p::GFP* marker in comparison to *ddi-1* mutants, which might reflect the fact that *png-1* and *skn-1a* are essential components of the proteasome surveillance pathway, while *ddi-1* has been shown to be dispensable [17]. Scale bar is 100 µm. **(C)** Gene structure of *ddi-1* and *png-1* showing the positions of mutant alleles used in this study. **(D)** Venn comparisons showing significant overlap between up-regulated genes in the transcriptome of *ddi-1(icb156)* & *skn-1a(mg570)* mutant, and bortezomib (BTZ)-treated animals with oomycete recognition response (ORR) (RF 13.8, *p* < 6.171e-50 for comparison with *ddi-1*; RF 13.6, *p* < 5.518e-101 for comparison with BTZ and RF 36.0, *p* < 1.249e-59 value for comparison with *skn-1a*). **(E)** *ddi-1(mg572), png-1(ok1654)* and *skn-1a(mg570)* mutant animals exhibit reduced susceptibility to infection by *M. humicola* as compared to WT *C. elegans* (n=60 per condition, performed in triplicates, *p*<0.001 based on log-rank test, representative graph shown).

Here, we show that, in contrast to previous studies demonstrating a protective role for SKN-1 in promoting resistance against bacterial pathogens, mutants in the SKN-1A-driven proteasome surveillance pathway result in constitutive expression of ORR and IPR and consequently exhibit resistance against infection by oomycetes and microsporidia. Rescue of *skn-1a* specifically in the epidermis or the intestine was sufficient to restore wild-type levels of infection by oomycetes and microsporidia respectively. Moreover, we report that the ORR/IPR genes induced in common in these programmes, are also regulated through the proteasome and can be induced in distinct tissues depending on the exact trigger. Notably, we show that by increasing proteasome function, there is a partial inhibition of oomycete-mediated induction of ORR markers, suggesting that the response to pathogens may directly involve stabilization of factors normally degraded by the proteasome. Therefore, our findings, together with other studies in flies and mammals [24–27], highlight how proteasome regulation keeps a range of innate immune responses in check in a tissue-specific manner.

## RESULTS

### Mutants in the proteasome surveillance pathway show constitutive activation of the ORR

We performed a chemical mutagenesis screen on animals carrying the *chil-27p::GFP* reporter, which is not expressed in wild-type animals under standard growth conditions, but is strongly induced upon recognition of oomycete pathogens [2, 28]. We obtained several independent mutants with constitutive epidermal expression of *chil-27p::GFP*, either in a graded manner with maximum GFP expression in the head region as previously reported [2], or strong expression throughout the body (Figure 1B). Three mutations were mapped to *ddi-1* and *png-1,* two genes known to play a role in the proteasome surveillance pathway through regulating SKN-1A [17, 29] (Figure 1A, C). To determine if the non-synonymous *ddi-1(icb156)* mutation isolated from the screen represented a gain or loss-of-function allele, we tested existing strong loss-of-function *ddi-1* mutants carrying either a deletion within the gene (*mg571*) or a mutation in the active site of the protein (*mg572*) [29]. In both cases, we found similar *chil-27p::GFP* induction (Figure S1A), confirming that *ddi-1(icb156)* represents a loss-of-function allele. Similarly, the recovery of a frameshift and a nonsense mutation in *png-1* suggested that constitutive *chil-27p::GFP* expression is likely attributed to loss-of-function of the gene, which was further confirmed using the independently derived *png-1(ok1654)* deletion allele [30] (Figure S1A).

As PNG-1 and DDI-1 are required for the activation of SKN-1A, we analyzed *chil-27p::GFP* expression upon loss of function of *skn-1a*. Here, we found constitutive expression throughout the body of the animal (Figure 1B). The response was also recapitulated by *skn-1* RNAi (Figure S1A), which is known to affect both *skn-1a* and *skn-1c* isoforms as the entire *skn-1c* sequence is shared by *skn-1a* (Figure S2A). Because PNG-1 is required specifically for activation of the SKN-1A isoform and SKN-1C does not undergo sequence editing [17], the involvement of SKN-1C in activating *chil-27p::GFP* seemed unlikely. However, to directly test this possibility, we performed *wdr-23* RNAi on *skn-1a(mg570)* mutant animals. WDR-23 is a WD40 protein known to specifically suppress SKN-1C function, inhibition which is released for example under oxidative stress to allow SKN-1C activation [31, 32]. We found that *wdr-23* RNAi did not suppress *chil-27p::GFP* induction in *skn-1a(mg570)* mutants (Figure S2B). This result suggests that activation of SKN-1C cannot rescue the constitutive expression of *chil-27p::GFP* in *skn-1a(mg570)* mutants, thus the constitutive response is likely due to loss of the *skn-1a* isoform.

Mutants in *skn-1a* have reduced proteasome subunit gene expression and, as a result, experience proteasome dysfunction [33]. Reduced proteasomal subunit gene expression may affect the induction of immune programmes through non-proteolytic roles that have recently been ascribed to proteasomal subunits, however induction in this case would not be affected by proteasome function impairment via bortezomib (BTZ) drug treatment [34]. We wanted to determine if impairment of proteasome function can induce the oomycete recognition response (ORR). We performed RNAseq analysis of *ddi-1(icb156)* mutants versus wild-type controls and compared the set of up-regulated genes with those previously reported for inhibition of proteasomal activity by BTZ drug treatment [35], oomycete recognition response [3] and *skn-1a* loss-of-function [35]. A significant overlap was identified between all these datasets (Figure 1D, S1B, and supplementary table S1). Wild-type animals treated with BTZ show induction of *chil-27* as previously described [15], and this induction is rapid at the *chil-27* mRNA level within 15 mins post exposure to the drug as opposed to 1 hour required upon exposure to oomycete extract (Figure S3), which further suggested a link between the induction of ORR and proteasome dysfunction. The physiological consequence of activated ORR was revealed by an infection assay where *ddi-1(mg572)*, *png-1(ok1654)* and *skn-1a(mg570)* mutants showed enhanced survival in the presence of the oomycete pathogen *Myzocytiopsis humicola* (Figure 1E). These findings are consistent with ORR induction in *skn-1a* mutants being protective against oomycete infection and associated with loss of proteasomal activity, as opposed to reduced proteasomal gene expression having an impact independent of proteasomal activity.

### Proteasome impairment in the epidermis is sufficient to induce the ORR

Induction of the ORR requires cross-tissue communication with chemosensory neurons sensing the pathogen and signalling to the epidermis where induction of *chil* genes takes place to combat the infection [3]. Having discovered that proteasome inhibition can activate the ORR, we wanted to determine whether this inhibition is required in a tissue-specific manner or not. To address this question, we performed tissue-specific rescue of *skn-1a(mg570)* mutants by expressing *skn-1a* under a *rab-3p* (pan-neuronal), *dpy-7p* (epidermal) or *vha-6p* (intestinal) promoter. We found that only the epidermal rescue of *skn-1a* repressed expression of *chil-27p::GFP* (Figure 2A). We further investigated this question by performing tissue-specific RNAi of the proteasomal subunit *rpt-5* [10, 36, 37], where we found that only epidermal RNAi of *rpt-5* was able to induce *chil-27p::GFP* expression (Figure 2B). We also tested the survival of tissue-specific rescued lines of *skn-1a* in the presence of *M. humicola* and found only epidermal overexpression of *skn-1a* to rescue the enhanced oomycete resistance phenotype (Figure 2C). These findings demonstrate that proteasome impairment, or rescue of proteasome surveillance function specifically in the epidermis, regulates the ORR and oomycete resistance.

**Figure 2:**
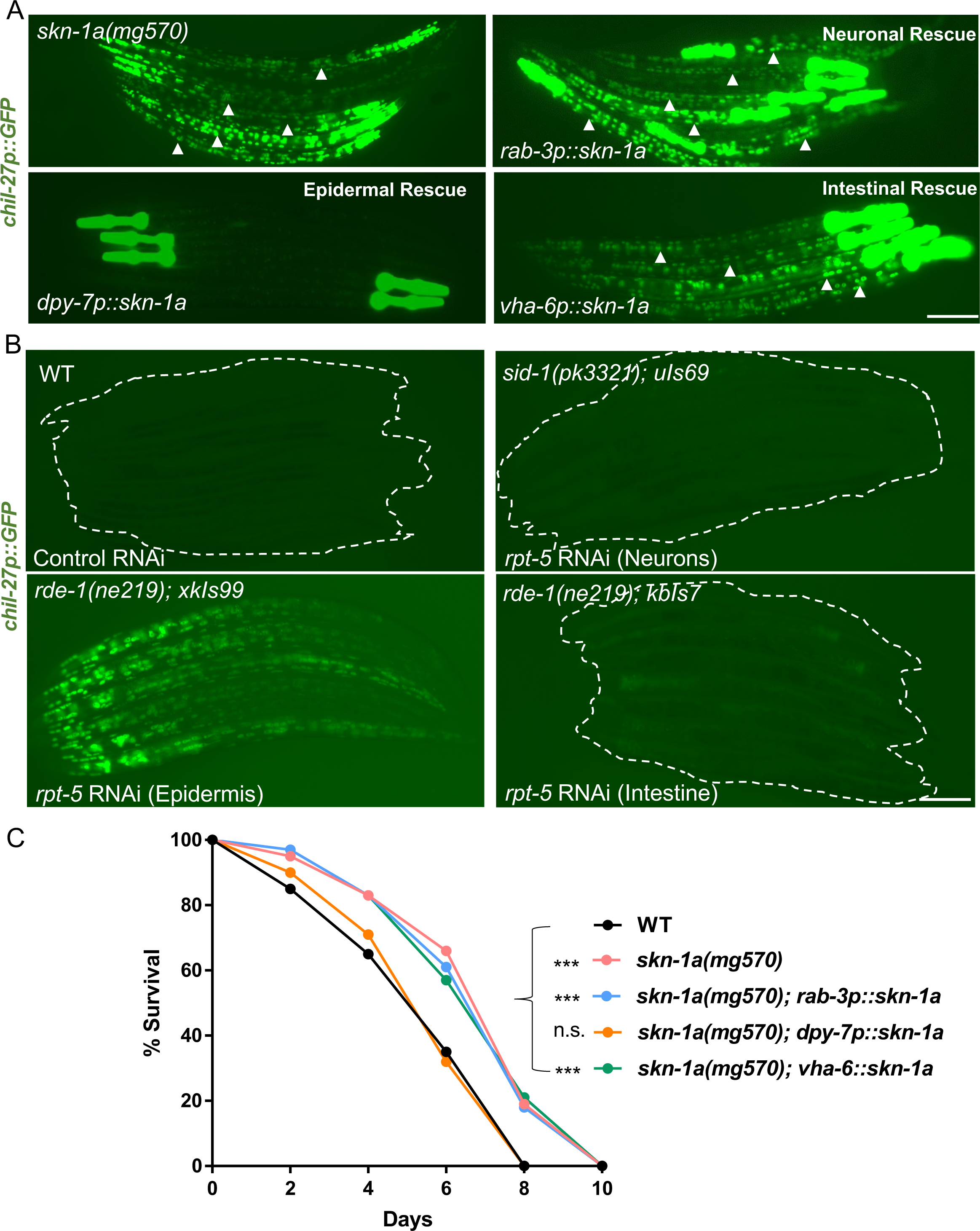
Proteasome impairment in the epidermis is sufficient to activate the ORR. **(A)** Epidermal (*dpy-7p*), neuronal (*rab-3p*) and intestinal (*vha-6p*) rescue of *skn-1a* function in *skn-1a(mg570).* Note loss of GFP puncta in the body (shown by arrowheads) corresponding to *chil-27p::GFP* expression specifically upon epidermal rescue. In all cases, *myo-2p::GFP* has been used as a co-injection marker which labels the pharynx. **(B)** Tissue-specific proteasome dysfunction induced by epidermal-specific, intestine-specific, and neuronal-enhanced RNAi of *rpt-5*. Scale bars in A and B are 100 µm. **(C)** Survival analysis of *skn-1a(mg570)* mutants with *skn-1a* function rescued in neurons (*rab-3p::skn-1a*), epidermis (*dpy-7p::skn-1a*) and intestine (*vha-6p::skn-1a*) (n=60 per condition, performed in triplicates, p<0.001 based on log-rank test, representative graph shown).

Previous work has led to the identification of other epidermal regulators of the ORR, namely the receptor tyrosine kinase OLD-1 [38] that is specific to oomycete recognition, and the PALS-22/PALS-25 antagonistic paralogs [15, 28], which regulate the immune response against both oomycetes and microsporidia. We thus asked whether activation of ORR upon proteasome dysfunction requires *old-1* or *pals-25.* Here, we performed *skn-1* RNAi on animals carrying a deletion in *old-1(ok1273)* or in *pals-25(jy81),* along with wild-type animals as control, and found no difference in *chil-27p::GFP* induction (Figure S4). These results suggest that epidermal proteasome dysfunction acts either downstream of OLD-1 and PALS-25-mediated signalling, or as a parallel trigger leading to the activation of ORR.

### Oomycete extract exposure does not cause broad proteasome dysfunction

Since epidermal proteasome dysfunction triggers the ORR, we tested whether impairment of proteasome function occurs upon exposure to oomycete extract. Proteasomal subunit expression is observed as a bounce-back response upon proteasome dysfunction [18]. Even though significant overlap was obtained between ORR, and genes upregulated upon proteasome dysfunction (Figure 1D), none of the proteasomal components were found to be induced as a part of the ORR (Figure 3A). For example, the induction of the *rpt-3p::GFP* reporter [29] was only observed upon BTZ treatment, but not upon treatment with oomycete extract (Figure 3B). Similarly, we did not see accumulation of ubiquitylated substrates reported by a *sur-5p::UbV-GFP* marker [39] upon extract exposure, while we did observe induction of the marker upon inhibition of proteasome activity by BTZ treatment (Figure 3C). These results suggest that oomycete extract exposure is unlikely to cause broad proteasome dysfunction in *C. elegans*.

**Figure 3:**
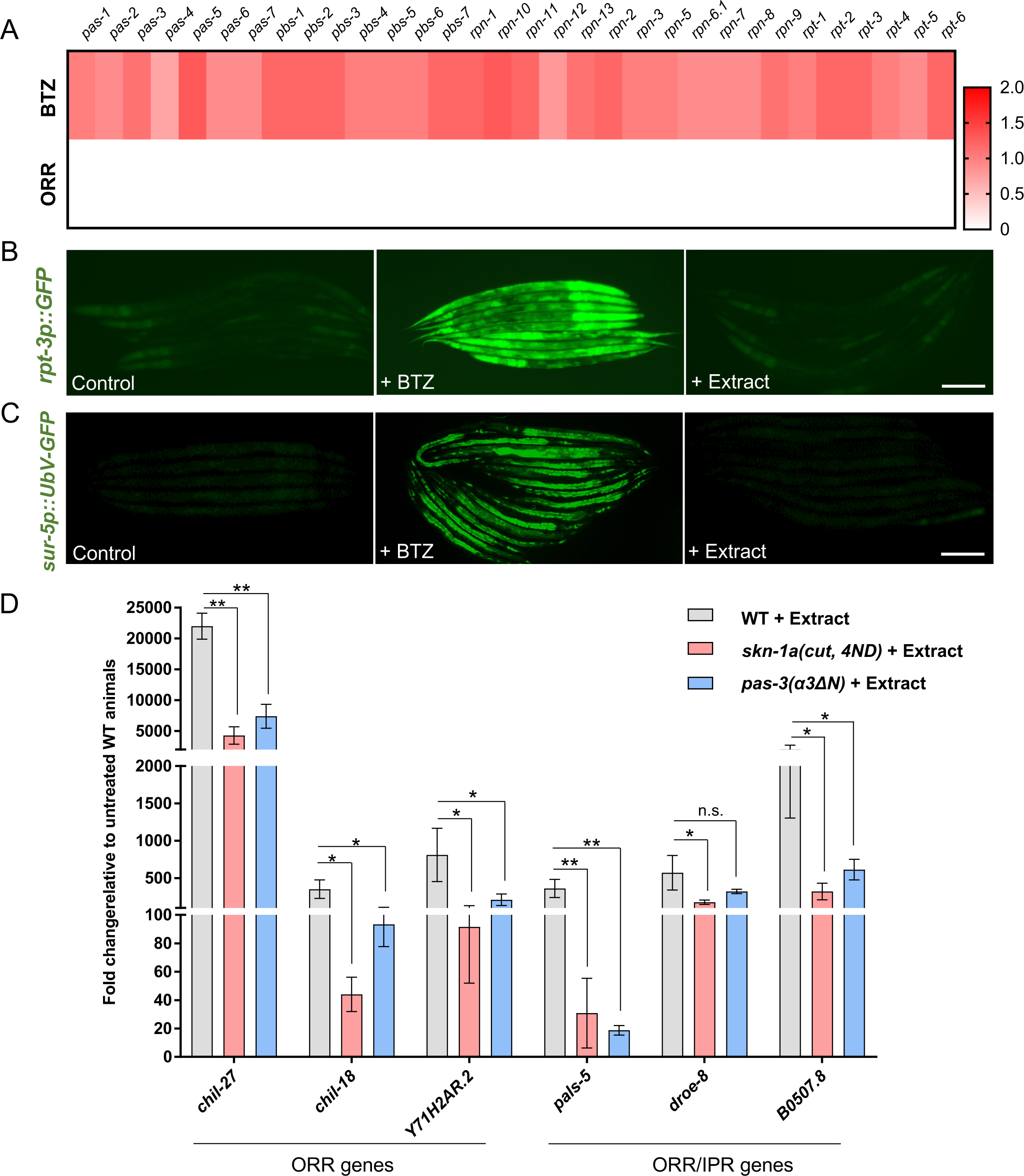
Oomycete extract exposure does not cause broad proteasome dysfunction, and activation of the proteasome inhibits induction of *chil-27* by oomycete extract. **(A)** Heat map showing log2 fold change in the expression of proteasome components upon treatment with oomycete extract as opposed to BTZ treatment. None of these genes are induced as part of the ORR and are shown in white. **(B)** Induction of *rpt-3p::GFP* is observed upon BTZ treatment, but not upon extract treatment. **(C)** Induction of *sur-5p::UbV-GFP* is observed upon BTZ treatment, but not upon extract treatment. Scale bar in panels B and C is 100 µm. **(D)** RT-qPCR showing reduced induction of genes specific to ORR or genes in the overlap between ORR and IPR upon extract treatment in animals with constitutive expression of the activated form of SKN-1A [*skn-1a(cut, 4ND)* in *skn-1a(mg570)*] or constitutive activation of the proteasome [*pas-3(α3ΔN)*] (\*\**p*<0.01, \*\*\*\**p*<0.0001 based on unpaired *t* test in comparison to extract-treated wild type).

To test the potential direct involvement of proteasomal regulation for activation of the ORR upon oomycete exposure, animals constitutively expressing proteasomal subunits through a constitutively activated SKN-1A *[skn-1a(cut, 4ND)]* [17] or animals having a hyperactive proteasome (*pas-3(α3ΔN)* [40] were treated with extract and expression of multiple ORR genes was analysed by RT-qPCR. We found partial but significant reduction in the expression of analysed ORR genes upon extract treatment in both cases (Figure 3D). These results suggest that pathogen-mediated induction of the ORR epidermal signalling pathway may partly require stabilization of a factor constitutively degraded by the proteasome under uninfected conditions.

### Impairment of the SKN-1A bounce-back response also leads to activation of resistance against intracellular intestinal pathogens

It is known that proteasome blockade by BTZ treatment can activate the IPR in *C. elegans* [15], which is the protective transcriptional program against intracellular intestinal pathogens like *Nematocida parisii* and the Orsay virus [4, 5]. To determine if mutants in the proteasome surveillance pathway also show constitutive activation of the IPR, we compared RNAseq datasets generated for *skn-1a* mutants and BTZ-treated animals with the IPR gene list. We found significant overlap between IPR and upregulated genes in *skn-1a* mutants (Figure 4A, B and Table S1). To investigate whether loss of *skn-1a* also leads to increased resistance against intestinal pathogens, we assayed the *skn-1a(mg570)* mutant for resistance against the intestinal pathogen *N. parisii*. Here, we found that *skn-1a(mg570)* mutants had increased resistance towards *N. parisii* (Figure 4C), which was rescued in this case specifically by intestinal (*vha-6p*) expression of *skn-1a* (Figure 4D). These results demonstrate that impairment of the SKN-1A proteasome surveillance pathway also induces resistance to intracellular pathogens of the intestine, and this impairment can be rescued specifically by SKN-1A function in the intestine.

**Figure 4:**
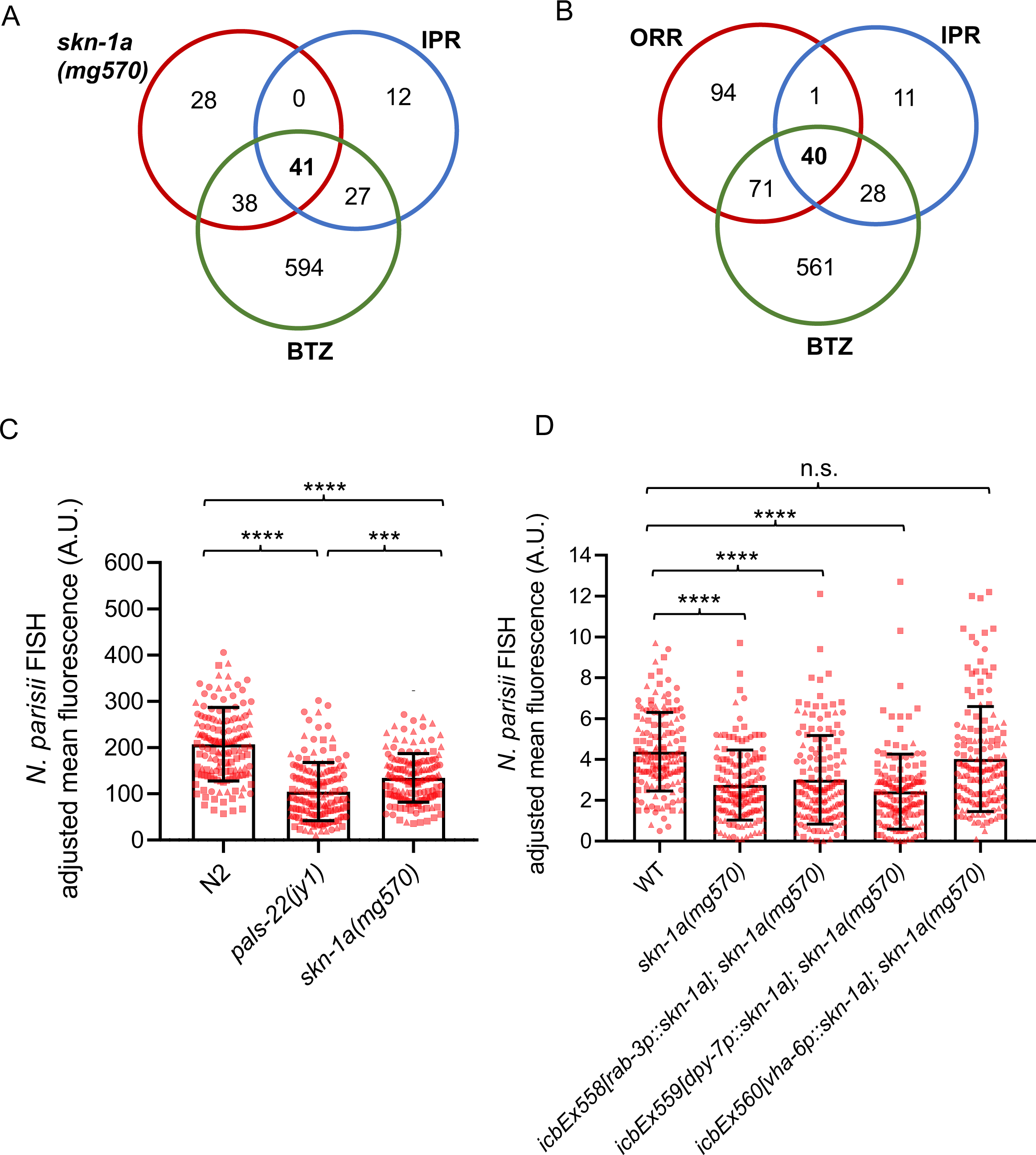
Proteasome impairment in the intestine leads to activation of the intracellular pathogen response (IPR). **(A)** Venn diagram showing significant overlap of IPR with up-regulated genes in *skn-1a* mutants (RF 84.4, *p* < 1.790e-72) and IPR with BTZ-treated animals (RF 21.4, *p* < 8.653e-84). **(B)** Venn diagram showing that overlap between ORR and IPR involves genes induced upon BTZ treatment. **(C)** *skn-1a(mg570)* mutants display increased resistance to *N. parisii* at 30 hpi relative to WT animals when infected at young adult. *pals-22(jy1)* mutant was used as a positive control which is already known to display increased resistance to *N. parisii*. **(D)** Intestinal rescue of *skn-1a,* but not epidermal or neuronal rescue, restore the resistance of *skn-1a(mg570)* mutants to *N. parisii* infection to WT levels. Kruskal-Wallis test with Dunn’s multiple comparisons test was used for statistical analysis (**** *p* < 0.0001, *** *p* < 0.001, ** *p* < 0.01. n = 150 animals per genotype).

We have previously reported that approximately half of IPR genes are also present in the ORR list [3], which was rather surprising given the distinct infection strategies and tissue tropism between oomycetes and microsporidia [41]. We hypothesised that the IPR and ORR programmes might share significant overlap because they both relate to regulation by the proteasome. Remarkably, we observed that almost all common ORR and IPR genes (40 out of 41 genes) were also regulated by BTZ treatment (Figure 4B and Table S1). We reasoned that these shared immune response genes may be induced in different tissues following an ORR or IPR trigger. To test this possibility, we made use of single molecule fluorescence *in situ* hybridization (smFISH) to determine in which tissue selected genes present in the overlap between ORR and IPR are induced, namely *pals-5, B0507.8, skr-3,* and *cul-6*. Animals carrying GFP-labelled epidermal nuclei (*dpy-7p::GFP-H2B*) were treated with oomycete extract to induce the ORR or were subjected to prolonged heat stress (24 h at 30°C) [5] as a proxy for inducing the IPR. Treatment with BTZ was performed to simultaneously activate both ORR and IPR. Consistent with our initial hypothesis, we found that all genes were induced specifically in the epidermis upon activation of ORR and in the intestine upon activation of IPR, while BTZ treatment led to induction in both tissues (Figure 5 and S5).

**Figure 5:**
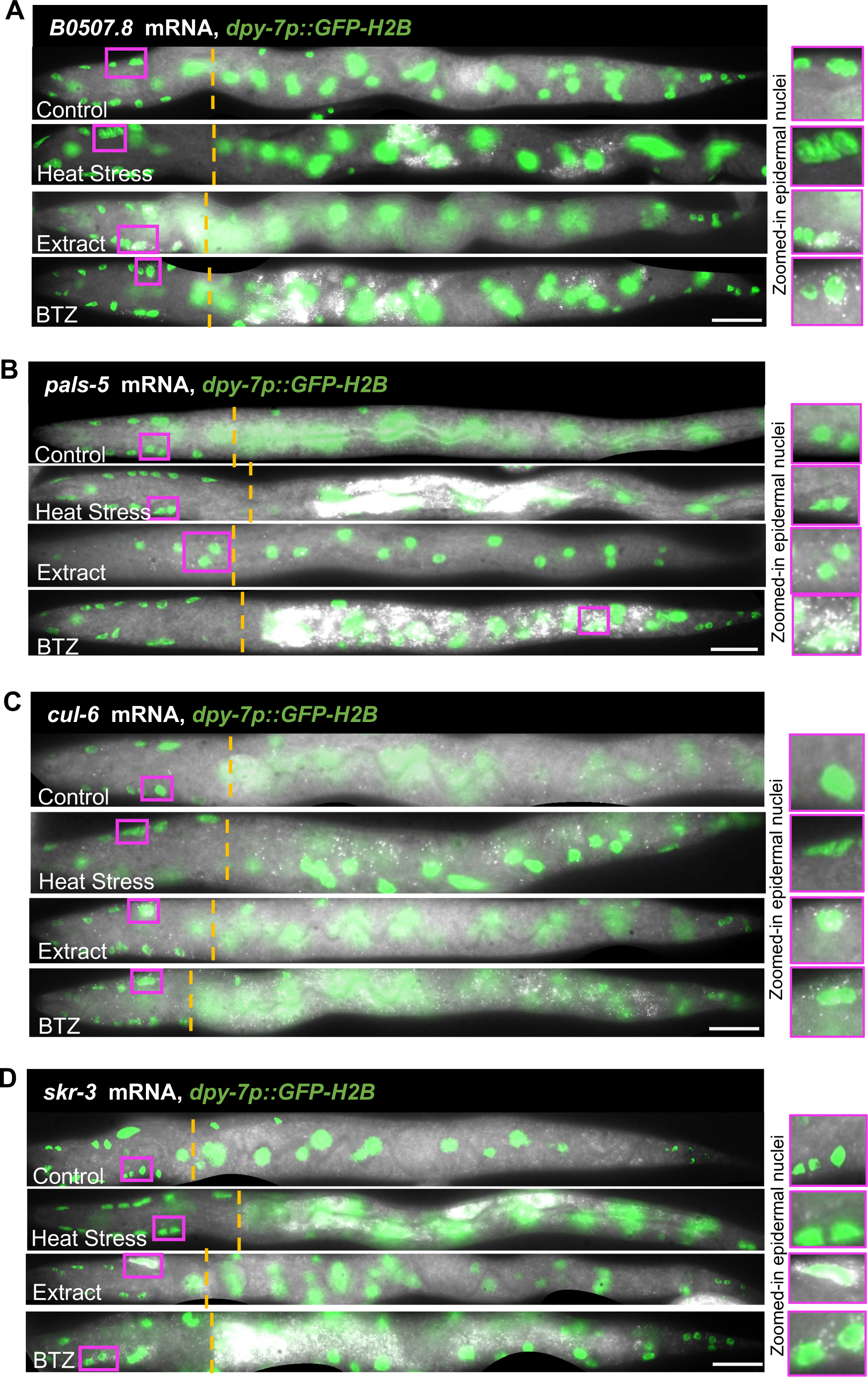
Common ORR and IPR genes can be induced in a tissue-specific manner. Sections of straightened L2 stage animals with head epidermal nuclei in focus showing mRNA distribution of some common ORR and IPR genes, namely *B0507.8* **(A)**, *pals-5* **(B)**, *cul-6* **(C)** and *skr-3* **(D)** by smFISH upon 4 hours extract treatment (ORR), prolonged heat stress at 30°C for 24 hours (IPR) and 1 hour of BTZ treatment (proteasome dysfunction). Epidermal nuclei are labelled in green with the *dpy-7p::GFP-H2B* marker. Co-localization of mRNA with green nuclei indicates epidermal expression as shown in zoomed-in panels (shown in magenta). Dashed orange line shows the beginning of the intestine extending towards the tail on the right. Scale bar is 10 µm.

Previous studies have linked *skn-1* to immunity against bacterial infection with *skn-1* loss-of-function mutants reported as being hypersusceptible to bacterial infection [21–23]. However, previous studies were performed with RNAi or mutants that affect multiple isoforms of *skn-1*, and so it was not clear whether *skn-1a* specifically was involved in bacterial immunity. We investigated this question by examining *skn-1a(mg570)* animals for increased susceptibility to *P. aeruginosa* (PA14) infection [42]. In this assay, *pals-22(icb89)* mutants and *skn-1* RNAi treated animals were used as positive controls, as both have been shown to result in enhanced susceptibility towards PA14 infection [15, 22]. The susceptibility of *skn-1a(mg570)* animals to PA14 was found to be comparable to wild-type animals, while both *skn-1* RNAi and *pals-22* mutants showed increased susceptibility as expected (Figure S6). The fact that *skn-1* RNAi targets both *a* and *c* isoforms, but *skn-1a(mg570)* animals are not hypersusceptible to PA14 suggests that requirement of SKN-1 to combat PA14 infection is more likely to be associated with the function of the SKN-1C isoform. Taken together, these results suggest that different *skn-1* isoform perturbations can lead to different host immunity outcomes in a pathogen-specific way.

## DISCUSSION

We demonstrate in this study that mutations in the SKN-1A proteasome surveillance pathway in *C. elegans* activate the ORR and IPR immune responses employed against distinct natural pathogens that infect through the epidermis or colonise the intestine. Our previous work indicated that blockade of the proteasome leads to induction of IPR genes, which are distinct from SKN-1-regulated genes [4, 15], but the connection between the SKN-1-regulated pathway and the ORR/IPR was not clear. Here we show that both loss of SKN-1A and impairment of proteasome function induce the genes in common to both the ORR/IPR. Furthermore, increasing proteasome function through constitutive proteasomal subunit gene expression or hyperactivity of the proteasome significantly inhibits oomycete extract-mediated *chil-27* gene induction. Taken together, we hypothesize that some proteasomally regulated factor(s) normally keep the ORR and IPR immune responses in check in the absence of infection, and upon exposure to specific pathogens such factors can be stabilized in the associated tissue triggering the rapid induction of the respective immune programmes (Figure 6). A similar phenomenon has been shown to regulate activation of NF-κB in *Drosophila,* where IMD is proteasomally degraded owing to permanent presence of Ub^K48^ linkages, which are lost upon bacterial infection thereby stabilizing the protein and activating the pathway [24, 25]. Likewise, Uba1-mediated proteasomal degradation of IRF3 in Zebrafish keeps activation of IFN signalling and activation of anti-viral immune response in check [43]. While our findings are consistent with a potential direct role for the proteasome in stabilising factors that may be necessary for ORR/IPR induction, we cannot rule out that the bounce-back response pathway may also act in parallel.

**Figure 6:**
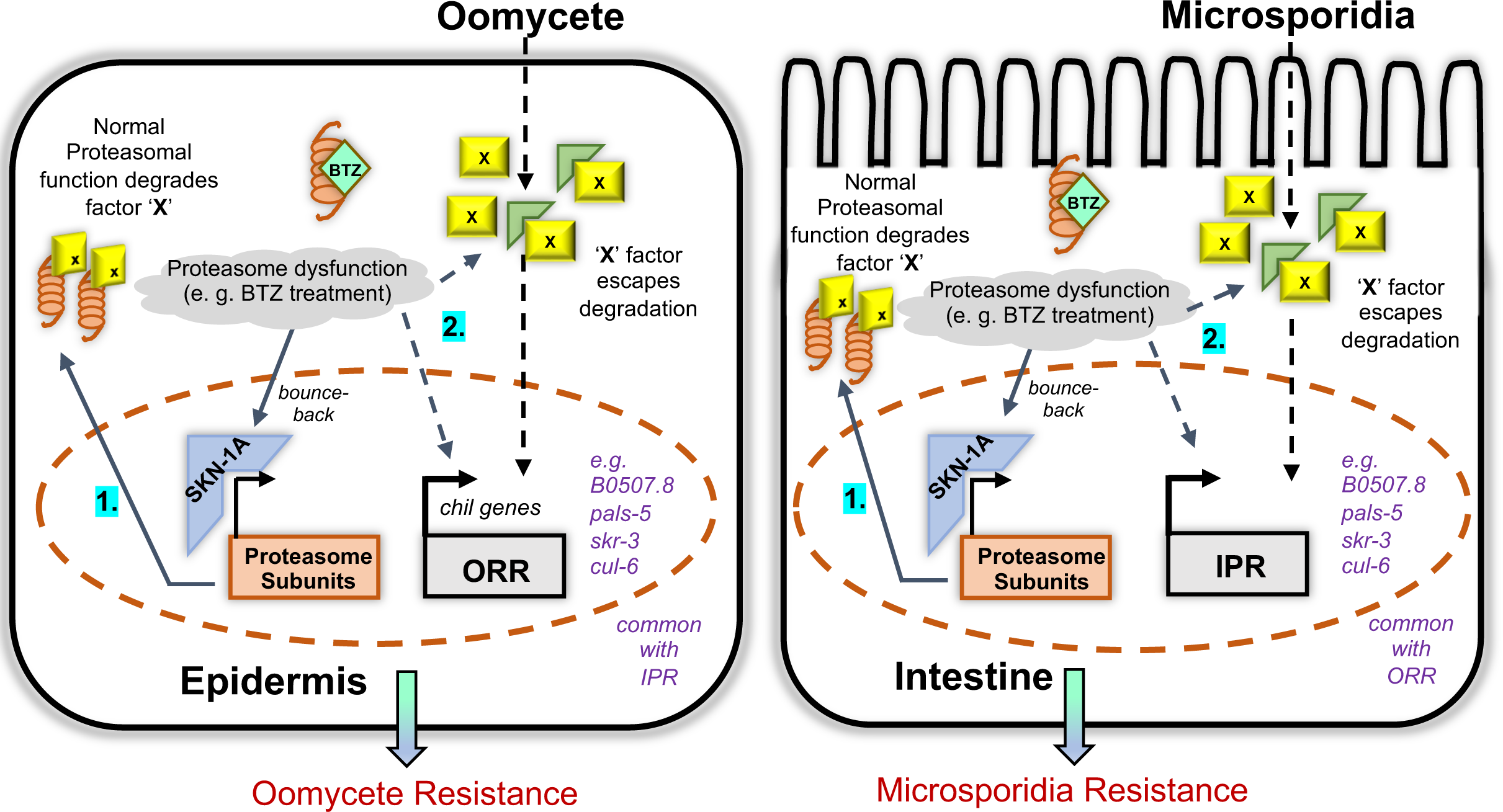
Model based on the findings of this study. The model highlights the interplay of SKN-1A mediated proteasomal gene expression and activation of ORR and IPR in the epidermis and the intestine respectively. **1.** Under normal conditions, proteasomal degradation of unknown positive regulators (denoted as X factor) keeps activation of immune responses in check. **2.** When proteasome dysfunction happens either by BTZ treatment or loss-of-function of *skn-1a*, X-factor escapes degradation and activates ORR and IPR in the epidermis and the intestine. Simultaneously, BTZ-mediated inhibition increases proteasomal gene expression as a bounce-back response. Such X factors could be directly involved in the tissue-specific signalling pathway activated upon oomycete or microsporidia exposure respectively in the epidermis and the intestine. Alternatively, they could regulate ORR and IPR gene induction in parallel to the pathogen-induced signalling pathway.

Unwanted activation of immune responses is often accompanied by trade-offs emphasizing the importance of keeping them in check [44]. For example, in *pals-22* loss-of-function mutants, constitutive activation of ORR/IPR provides them protection from oomycetes and microsporidia [3]. However, *pals-22* mutants also exhibit slow development, reduced lifespan, and increased pathogen load upon exposure to *P. aeruginosa* [15]. Likewise, *pals-17* loss-of-function mutants exhibit constitutive activation of IPR and immunity to intracellular pathogens, but impaired development and reproduction [45]. While *skn-1a(mg570)* mutants develop normally like wild-type *C. elegans*, they too exhibit increased age-associated protein aggregation and consequently have a reduced lifespan [33], so similar trade-offs may also come into play in this case and involve proteotoxicity-driven defects that may exacerbate upon stressful conditions. Likewise, human patients with proteasome-associated autoinflammatory syndromes (PRAAS) show hyperactivation of IFN signalling, but often suffer with severe neurodevelopmental anomalies and skeletal defects [26, 27, 46].

Our study also highlights the importance of isoform-specific functions for *skn-1.* While *skn-1b* is neuronally expressed and regulates satiety and metabolic homeostasis [47], *skn-1a* and *skn-1c* are shown to perform antagonistic immunomodulatory functions in a pathogen-specific way. Previous studies that reported a requirement of *skn-1* for bacterial resistance used either *skn-1* RNAi or *skn-1* mutant alleles which affected both *a* and *c* isoforms [22, 23]. However, we now show that *skn-1a* mutants show enhanced resistance to infection by eukaryotic natural pathogens and do not show enhanced susceptibility to PA14, which suggests that bacterial immunity is likely to be driven by SKN-1C. This result also aligns with the fact that just like NRF2 in humans, SKN-1C regulates response to oxidative stress in *C. elegans* [20], and bacterial pathogens have been shown to trigger ROS production in the host [21].

While proteasome-mediated suppression of inflammatory/immune responses might be an evolutionarily conserved phenomenon, the exact mechanisms of how immune responses are kept in check are likely to differ. For example, activation of type-I interferon signalling in mammals as a consequence of proteasome dysfunction has been attributed to ER stress and activation of the unfolded protein response (UPR) [26, 48, 49]. This is unlikely to be the case in the context of oomycete recognition as we have previously shown that introducing ER stress does not induce *chil-27p::GFP* [2]. Similarly, markers of proteotoxic stress are not induced as a part of the ORR/IPR programs, which makes them distinct from other stress-induced responses. Whether proteasomal regulation of the same factors keeps immune responses in check in the *C. elegans* epidermis and intestine is currently unknown. In the epidermis, while there is evidence that protein levels for the receptor tyrosine kinase OLD-1, which is fully required for mounting the ORR, are regulated by the proteasome [38], *old-1* knockouts are still able to mount the ORR upon *skn-1a* perturbation ruling out OLD-1 as the main driving factor. In the intestine, ZIP-1 is a plausible candidate because it is degraded by the proteasome under basal conditions and has been shown to be required for induction of a subset of IPR genes [50]. Future work will elucidate the exact proteasomally-regulated triggers of ORR/IPR and address whether these are shared or not between different tissues in *C. elegans*.

## MATERIALS AND METHODS

### *C. elegans* strains and pathogen maintenance

*C. elegans* strains were cultured on NGM plates seeded with *E. coli* OP50 at 20°C under standard conditions as previously described [51]. The list of all the strains used in this study is provided in supplementary table S2. We refer to *skn-1(mg570)* mutants throughout the manuscript as *skn-1a(mg570)* to stress that only isoform *a* is affected. Both *M. humicola* and *N. parisii* were maintained as described previously [2, 41].

### EMS mutagenesis

L4 stage WT animals carrying the *chil-27p::GFP* reporter (*icbIs4*) were mutagenized with 24 mM ethyl methanesulfonate (EMS) (Sigma) in 4 mL M9, for 4 hours with intermittent mixing by inversion. Worm pellets were then washed 10 times with 15 mL M9 to completely get rid of EMS from the suspension and worms were plated onto 90 mm NGM plate seeded with *E. coli* OP50. After 24 h, 300 adults were randomly picked and divided into 30 90 mm plates carrying 10 animals in each. After 72 hours at 20°C, all F1 gravid adults in each plate were individually bleached and respective pool of F2 embryos were collected onto new plates. Around 60,000 haploid genomes were screened to identify animals showing constitutive activation of *chil-27p::GFP*. The causative mutations were mapped to *ddi-1* and *png-1* by crossing to the highly polymorphic CB4856 strain, and 15-25 F2 recombinants were pooled in equal proportion to obtain genomic DNA for whole genome sequencing of each independent mutant, performed by BGI (Hongkong). WGS data was analysed using the CloudMap Hawaiian variant mapping pipeline to identify causative mutations [52]

### Oomycete infection assays

Six *M. humicola* infected (dead) animals were added to the lawn of *E. coli* OP50 on three NGM plates and 30 live L4 animals for each genotype were transferred individually to each plate (n=90 per condition per repeat). Dead animals with visible sporangia were scored every 48 hours and live animals were transferred to a new NGM plate with six *M. humicola* infected (dead) animals. Dead animals without evidence of infection or animals missing from the plate were censored. Infection assays were performed in triplicates at 20°C on 30 mm NGM plates seeded with 100 µl *E. coli* OP50.

### *P. aeruginosa* PA14 infection assays

100µL of overnight grown PA14 culture in LB was spread on a 55mm NGM plate containing 10µg/mL FuDR. The plates were incubated at 37°C for 24 h. 90 Day 1 adults grown on NGM-OP50 or NGM-HT115/*skn-1* RNAi in the presence of FuDR were transferred to 3 plates each containing 30 animals (in triplicates), and incubated at 25°C. The number of animals alive were counted every 24h and dead animals defined as unresponsive to touch were removed from the plate. The experiment was continued until all animals on the plate were dead.

### Microsporidia *N. parisii* infection assays

600 synchronized L1 stage worms for each genotype were plated to NGM + *E. coli* OP50-1 and grown to the young adult life stage at 23°C (48 h for all stains except for *pals-22(jy1)* mutants, which are developmentally delayed [5] and therefore were grown for 52 h). The 600 young adults were washed off the plate with M9 + 0.1% Tween-20 (M9-T), pelleted, and the supernatant was removed. The worms were then mixed with 2 million *N. parisii* spores (strain ERTm1), 50 µl of 10X concentrated *E. coli* OP50-1, and M9 to a final volume of 300 µl. The infection mix was top-plated over the entire surface of a 6-cm NGM plate, dried at room temperature, and transferred to 25°C for 3 h. Following the 3 h pulse infection, the worms were collected from the infection plate with M9-T into a 1.5 ml microcentrifuge tube, pelleted, and then washed three additional times with 1 ml of M9-T. The infected worms were then transferred to an NGM + *E. coli* OP50-1 plate without *N. parisii* spores and returned to 25°C for an additional 27 h. Infected animals were then fixed in 100% acetone and incubated at 46°C overnight with FISH probes conjugated to the red Cal Fluor 610 fluorophore that hybridize to *N. parisii* ribosomal RNA (Biosearch Technologies). Samples were analyzed for *N. parisii* meronts [53] using an ImageXpress Nano plate reader using the 4x objective (Molecular Devices, LLC). The worm area was traced using FIJI software and the average red fluorescence intensity of each worm was quantified with the background fluorescence of the well subtracted. 50 animals per genotype were quantified for each experimental replicate, and three independent infection experiments were performed.

For the tissue-specific *skn-1a* rescue strains MBA1660, MBA1661, and MBA1662 approximately 100 *bus-1p::GFP*-expressing worms were manually picked onto *N. parisii* infection assay plates. For consistency, approximately 100 adults for WT and *skn-1a(mg570)* controls were also picked in the same manner and pulse infection was performed as described above. To remove any potential bias, the order in which strains were picked was randomized from assay to assay, and the 30 hpi endpoints were staggered to account for picking time. Fixation and FISH hybridization were performed as described above. For quantification, fixed animals for each strain were mounted to a 5% agarose pad on a microscope slide and imaged on a Zeiss AxioImager M1 compound microscope using a 2.5x objective. The worm area was traced using FIJI software and the average red fluorescence intensity of each worm was quantified with the background fluorescence of the slide subtracted. In some samples, we observed dim red autofluorescence from embryos inside the adult gonad. Thus, this region of the worm was omitted in all samples when tracing the worm area. Analysis with the ImageXpress plate reader and Zeiss AxioImager produced different arbitrary values for *N. parisii* FISH pixel intensity. However, the fold difference in infection between WT animals and *skn-1a(mg570)* mutants was similar (WT animals are ∼1.5 fold more infected than *skn-1a(mg570)*) by both imaging and quantification methods. 50 animals per genotype were quantified for each experimental replicate, and three independent infection experiments were performed.

### RNAseq

For transcriptome analysis, synchronised L4 stage animals were collected in triplicates for RNA extraction using TRIzol (Invitrogen) and isopropanol/ethanol precipitation. RNA sequencing was completed by BGI (Hongkong). Kallisto [54] was used for alignments with the WS283 transcriptome from Wormbase. Count analysis was performed using Sleuth [55] along with a Wald Test to calculate log_2_fold changes. All RNAseq data files are publicly available from the NCBI GEO database under the accession number GSE241087.

### RT-qPCR

Synchronized L4 stage animals with 4-hour extract treatment or without were collected in TRIzol (Invitrogen) followed by RNA extraction using isopropanol/ethanol precipitation. RNA was quantified using NanoDrop (Thermo Scientific) and its quality was analyzed by gel electrophoresis. cDNA was synthesized using 2 µg RNA with Superscript IV (Invitrogen) and Oligo(dT) primers as per manufacturer’s instructions. Real-time PCR was performed using qPCR primer pairs listed in Table S2 and LightCycler480 SYBR Green I Master Mix (Roche) in a LightCycler480 instrument; and Ct values were derived using the LightCycler480 software and second derivative maximum method. All experiments were performed in biological triplicates and changes in expression were calculated via the 2-ΔΔCt method.

### Microscopy

Animals to be imaged were picked into a drop (5-7 µL) of M9 containing sodium azide (50 µM) on an agarose (1%) pad made on a glass slide. Coverslip was added once the animals were paralyzed, and the pad was almost dry. *chil-27p::GFP/col-12p::mCherry* expression was observed on Zeiss Axio Zoom V16 microscope and images were taken using the associated ZEN Microscopy software. For *rpt-3p::GFP* and *sur-5::UbV-GFP*, expression was observed on Zeiss Compound microscope (AxioScope A1) at 40x magnification and images were taken using OCULAR.

### Molecular cloning and transgenesis

To generate constructs for tissue-specific rescue of *skn-1a(mg570)* mutant, *skn-1a* fragment was amplified from N2 cDNA using primers skn-1a_fullF and skn-1a_fullR. Promoter fragments namely, *rab-3p* and *vha-6p* were amplified from N2 genomic DNA using primer pairs BJ97-pRab-3Fwd, rab-3rev-skn1, and bj97 vha-6 Fwd, vha-6 Rev skn-1a respectively. The terminator sequence from 3’UTR of *unc-54* was PCR amplified from N2 genomic DNA using primers unc-54-F-skn1a and BJ36_unc-54terR. Plasmids for neuronal and intestinal rescue were assembled with specific promoter, *skn-1a* and *unc-54* 3’UTR fragments ligated into SpeI (FastDigest)-digested pBJ97 by Gibson cloning. To make the epidermal rescue plasmid, *skn-1a* was PCR amplified from N2 cDNA using primers dpy-7_skn-1F and dpy-7_skn-1R and was assembled into PmeI(FastDigest)–digested pIR6 by Gibson cloning. All constructs were individually injected into *skn-1a(mg570)* at 5 ng/µL with 25 ng/µL of pRJM163 (*bus-1p::GFP*) as the co-injection marker and 80 ng/µL of pBJ36 as carrier DNA. The transgenic strains thus created were used for microsporidia pathogen load assays. Similarly, all constructs were individually also injected in the strain MBA1055, which is *skn-1a(mg570)* mutant with *chil-27p::GFP* reporter in the background. For these injections, 5 ng/µL of rescue plasmid with 5 ng/µL of *myo-2p::GFP* as co-injection marker and 100 ng/µL of pBJ36 as carrier DNA was used. The transgenic strains thus created were analysed for expression of *chil-27p::GFP* in *skn-1a(mg570)* mutants.

### RNAi

RNAi by feeding was used as the means to induce gene knockdown in *C. elegans* as previously described [56]. The RNAi clone for *skn-1* was obtained from the ORFeome Library (Horizon Discovery) [57] and for *wdr-23* and *rpt-5* were obtained from the Ahringer Library (Source BioScience) [58]. Both the clones were confirmed by sequencing prior to use. Embryos collected by bleaching gravid adults were added onto NGM plates seeded with *E. coli* HT115 and contained 1 mM IPTG (Sigma) to induce expression of dsRNA. After 72 hours of incubation at 20°C, animals at L4/adult stage were scored for *chil-27p::GFP* expression (n>50). The strains used for tissue-specific RNAi were generated by introducing the *chil-27p::GFP* reporter in previously described strains listed in Table S2 with their associated references.

### smFISH

For inducing the ORR, synchronized L2 stage animals were treated with oomycete extract for 4 hours. For inducing IPR, animals were subjected to prolonged heat stress by incubating synchronized L1s at 30°C for 24 hours. For causing proteasome dysfunction, synchronized L2 stage animals were treated with 20 µM BTZ treatment for 15 min or 2 hours. Following requisite treatment along with relevant controls, animals were fixed with 4% formaldehyde (Sigma-Aldrich) in 1x PBS (Ambion) for 45 min and were permeabilised with 70% ethanol for 24 hours. Hybridization was performed at 30°C for 16 hours. List of oligos included in all the probes can be found in Table S2. Imaging was performed in an inverted and fully motorised epifluorescence microscope (Nikon Ti-eclipse) with an iKon M DU-934 CCD camera (Andor) controlled via theNIS-Elements software (Nikon) using the 100x objective. Detailed protocol can be found in [59].

## ACKNOWLEDGEMENTS

We thank Domenica Ippolito and Kenneth Liu for comments on the manuscript. Some strains were provided by the CGC, which is funded by NIH Office of Research Infrastructure Programs (P40 OD010440). We thank Gary Ruvkun, Thorsten Hoppe, and David Smith for strains. This work was supported by the Wellcome Trust [219448/Z/19/Z] and the BBSRC [BB/X001865/1].

## SUPPLEMENTARY FIGURE LEGENDS

**Figure S1:**
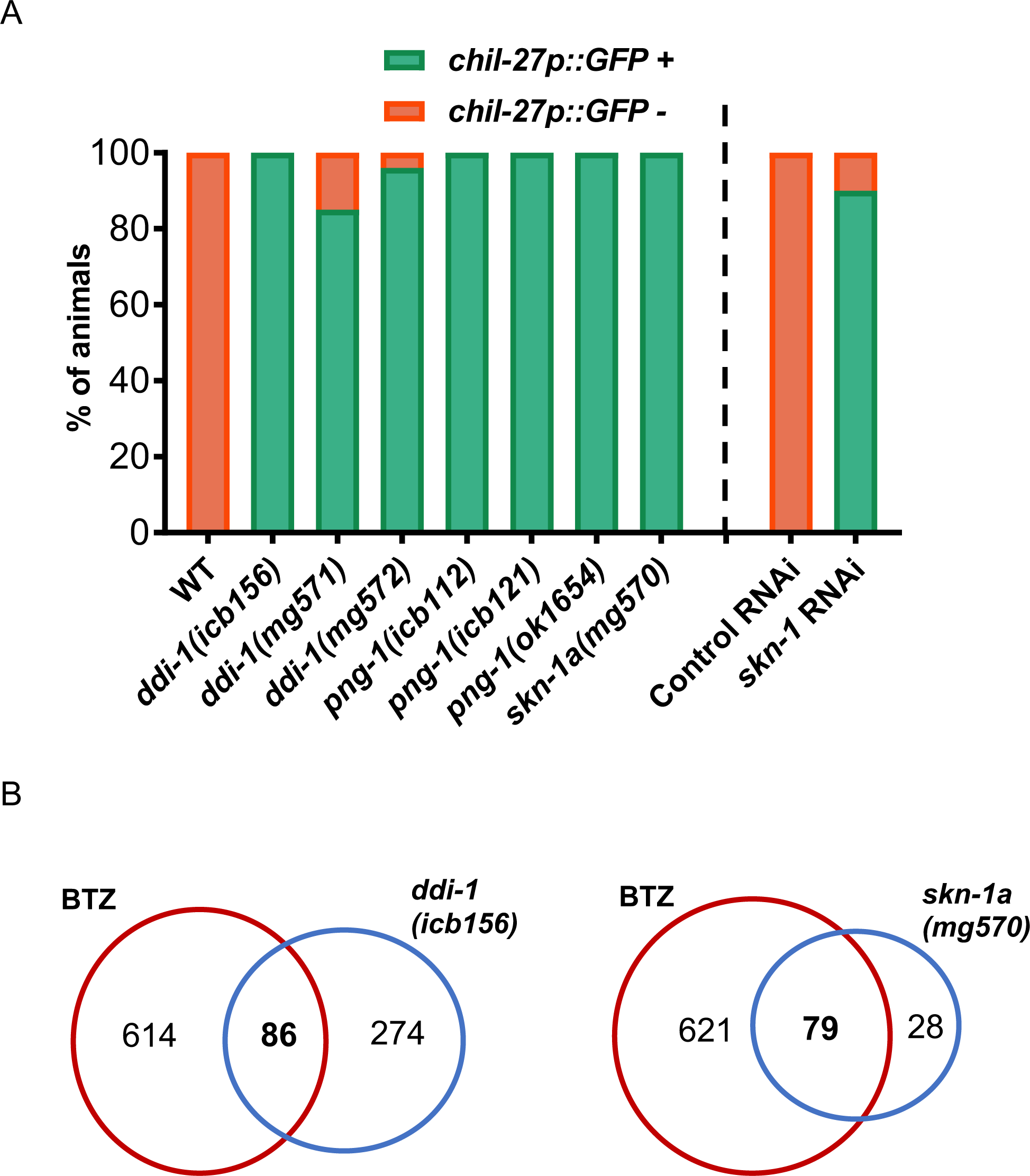
Additional alleles of proteasome surveillance mutants and *skn-1* RNAi show constitutive *chil-27p::GFP* expression. **(A)** Quantification of induction of *chil-27p::GFP* in the mutants obtained from the EMS screen, *ddi-1(icb156), png-1(icb112), png-1(icb121)* along with active-site mutant of *ddi-1(mg572),* in-frame gene deletion of *ddi-1(mg571), png-1(ok1654), skn-1a(mg570)* and upon *skn-1* RNAi (n>50, \*\*\*\**p* value < 0.0001, *** *p* value < 0.001 based on chi-square test). **(B)** Venn comparisons showing significant overlap between up-regulated genes in the transcriptome of *ddi-1(icb156)* and *skn-1a(mg570)* mutant with bortezomib (BTZ)-treated animals (RF 6.0, p < 4.347e-43 and RF 18.6, p < 4.453e-88 respectively).

**Figure S2:**
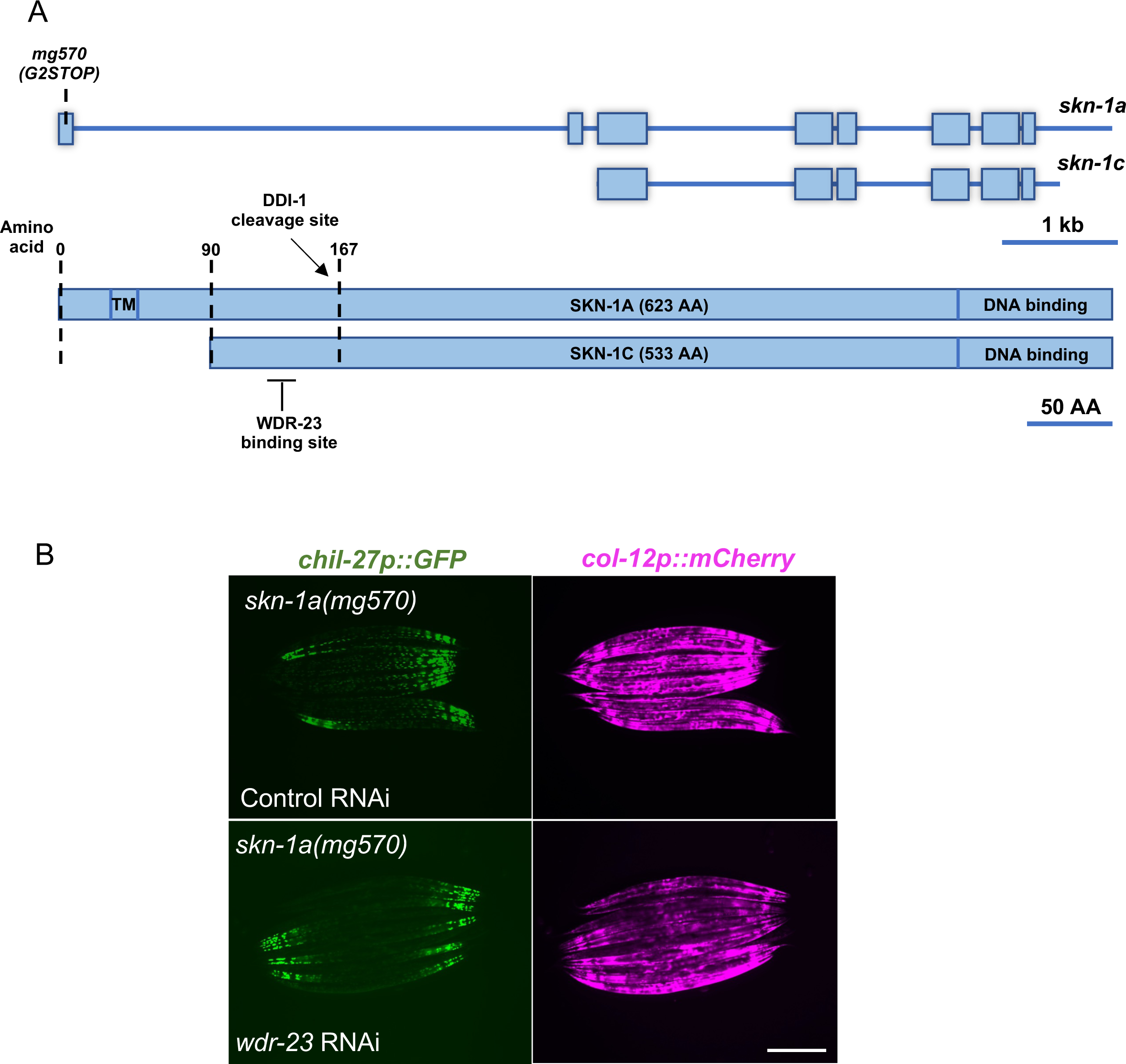
*wdr-23* RNAi on *skn-1a(mg570)* mutants. **(A)** Gene structure of *skn-1a* and *skn-1c* isoforms with protein domain organization. **(B)** Day 1 adults showing constitutive *chil-27p::GFP* expression upon *wdr-23* RNAi and control RNAi in *skn-1a(mg570)* mutants. Scale bar is 100 µm.

**Figure S3:**
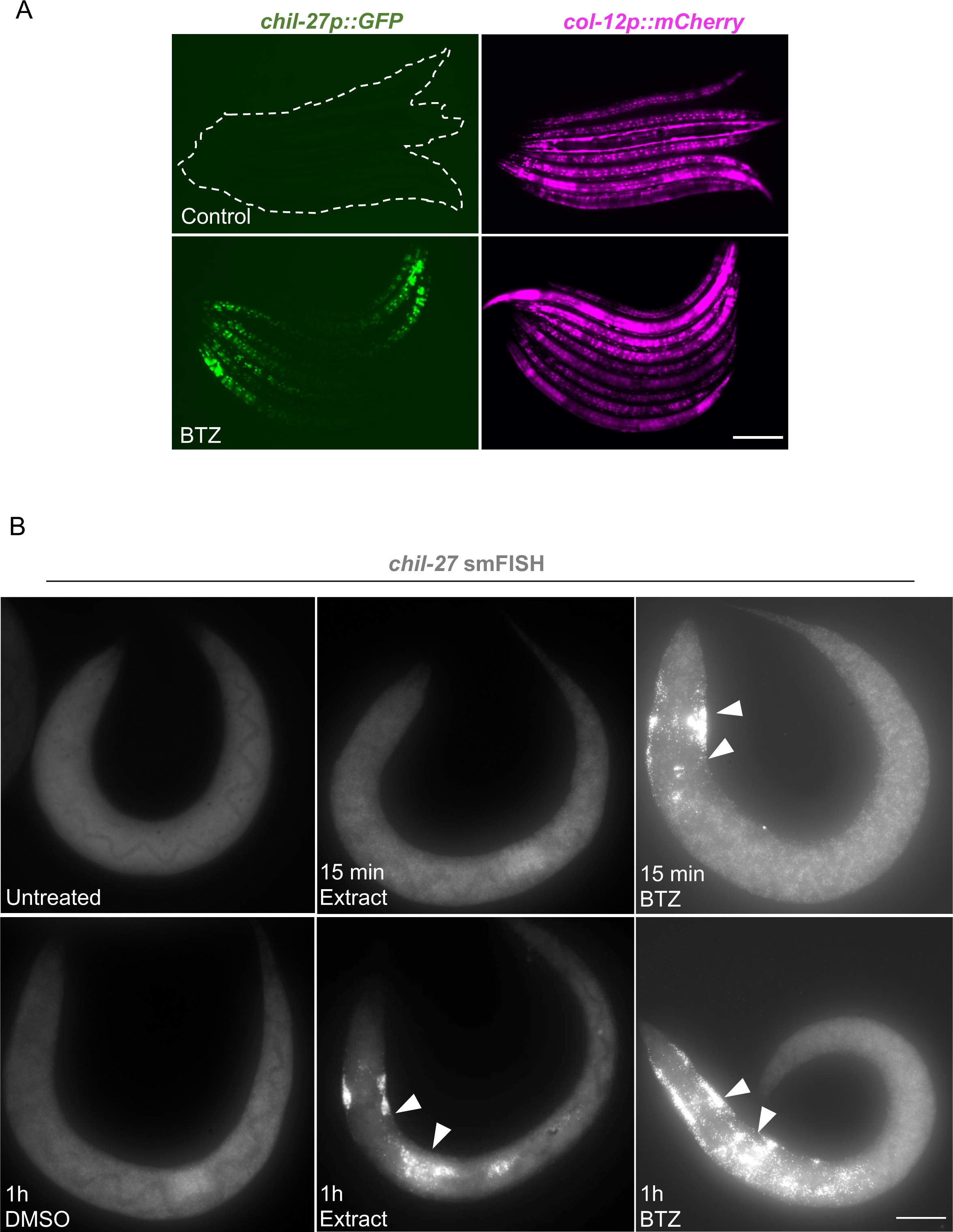
Bortezomib (BTZ) treatment induces *chil-27* expression. **(A)** WT L4 stage *C. elegans* showing *chil-27p::GFP* expression upon 20 µM BTZ treatment for 24 hours. **(B)** Z-stack of L2 stage WT animals showing expression of *chil-27* mRNA upon 15 min and 1 hour post extract and BTZ treatment. Scale bar is 100 µm in (A) and 10 µm in (B).

**Figure S4:**
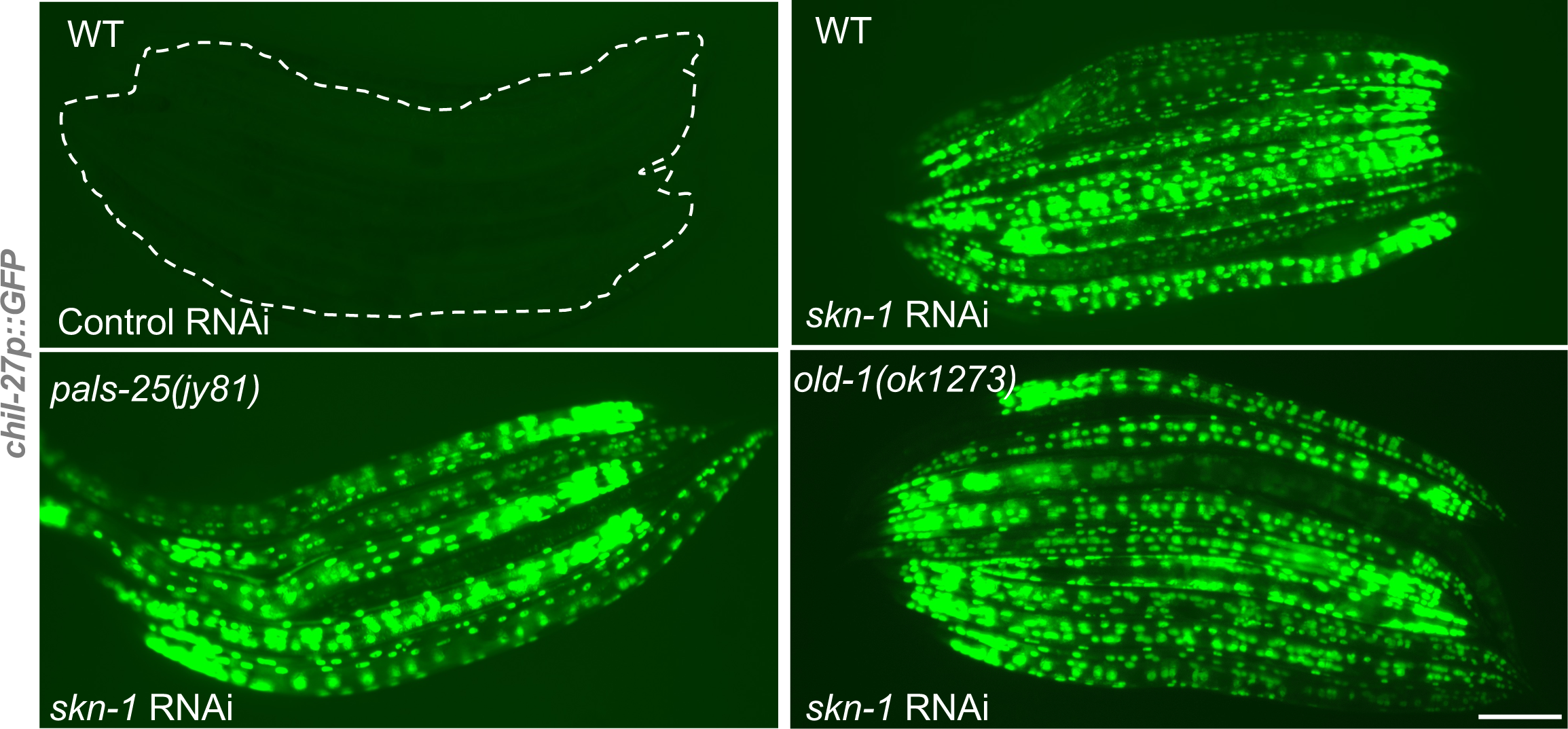
Epistasis analysis of *skn-1* and other regulators of the ORR. *skn-1* RNAi induces *chil-27p::GFP* expression in *pals-25(jy81)* and *old-1(ok1273)* mutant animals. Scale bar for all panels is 100 µm, and n>50 per condition.

**Figure S5:**
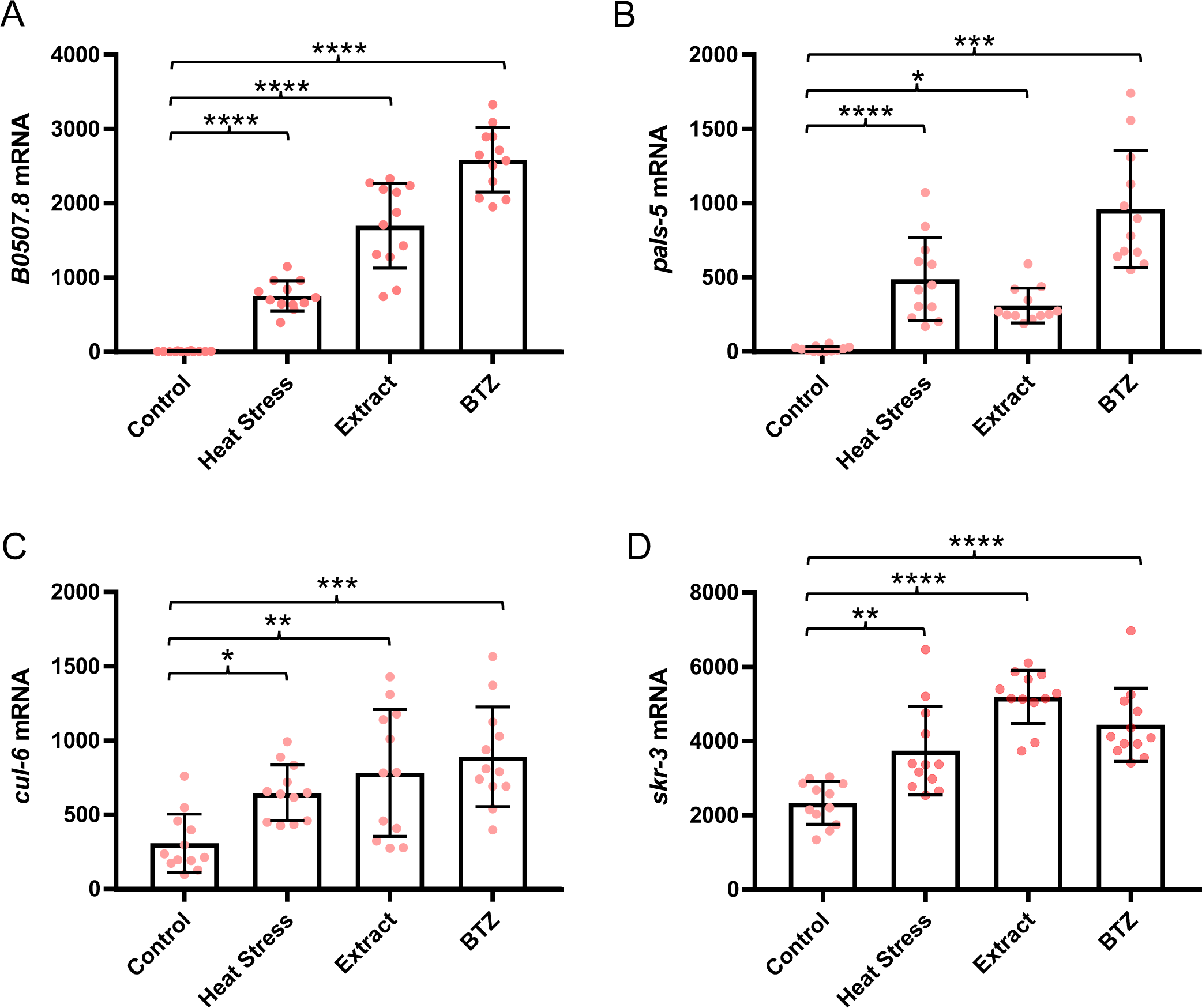
Quantification of expression of common ORR and IPR genes. mRNA quantification for four genes namely *B0507.8* **(A)**, *pals-5* **(B)**, *cul-6* **(C)**, and *skr-3* **(D)** quantified by smFISH in L2-stage animals subjected to prolonged heat stress (30°C for 24 hours), extract treatment (4 hours) and 20 µM BTZ treatment (1 hour). (n=12 in each condition and one-way ANOVA and Tukey’s multiple comparison test was used to assess statistical significance, \**p*<0.05 \*\**p*<0.01, \*\*\**p*<0.001, \*\*\*\**p*<0.0001).

**Figure S6:**
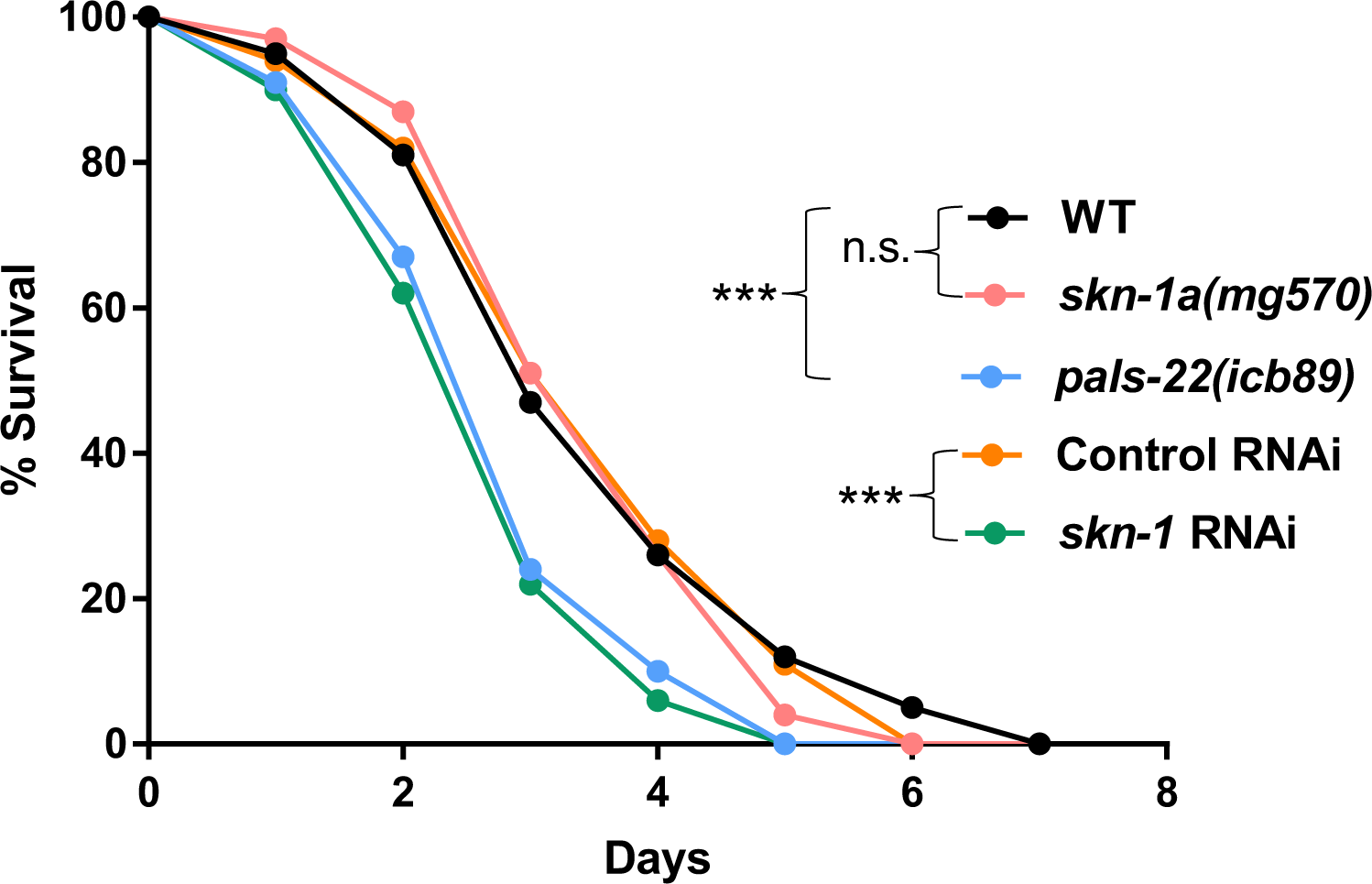
*skn-1a(mg570)* mutants do not show hyper-susceptibility to *P. aeruginosa* PA14. Survival analysis of adult *C. elegans* on PA14 where bacteria was spread on the plate (n=90 per condition, performed in triplicates, *p*<0.001 based on log-rank test, representative graph shown).

## SUPPLEMENTARY TABLE LEGENDS

**Table S1:** Differentially expressed genes in *ddi-1(icb156)* mutants as compared to WT animals; along with datasets used to make Venn diagrams, and the list of overlapping genes in different conditions.

**Table S2:** List of strains and oligos used in the study.

## REFERENCES

1. Schulenburg H, Felix MA. The natural biotic environment of *Caenorhabditis elegans*. Genetics. 2017;206(1):55–86. doi: 10.1534/genetics.116.195511. PubMed PMID: 28476862; PubMed Central PMCID: PMCPMC5419493.

2. Osman GA, Fasseas MK, Koneru SL, Essmann CL, Kyrou K, Srinivasan MA, et al. Natural infection of *C. elegans* by an oomycete reveals a new pathogen-specific immune response. Curr Biol. 2018;28(4):640–8 e5. Epub 2018/02/06. doi: 10.1016/j.cub.2018.01.029. PubMed PMID: 29398216.

3. Fasseas MK, Grover M, Drury F, Essmann CL, Kaulich E, Schafer WR, et al. Chemosensory neurons modulate the response to oomycete recognition in *Caenorhabditis elegans*. Cell Rep. 2021;34(2):108604. Epub 2021/01/14. doi: 10.1016/j.celrep.2020.108604. PubMed PMID: 33440164; PubMed Central PMCID: PMCPMC7809619.

4. Bakowski MA, Desjardins CA, Smelkinson MG, Dunbar TL, Lopez-Moyado IF, Rifkin SA, et al. Ubiquitin-mediated response to microsporidia and virus infection in *C. elegans*. PLoS Pathog. 2014;10(6):e1004200. Epub 20140619. doi: 10.1371/journal.ppat.1004200. PubMed PMID: 24945527; PubMed Central PMCID: PMCPMC4063957.

5. Reddy KC, Dror T, Sowa JN, Panek J, Chen K, Lim ES, et al. An intracellular pathogen response pathway promotes proteostasis in *C. elegans*. Curr Biol. 2017;27(22):3544–53 e5. Epub 2017/11/07. doi: 10.1016/j.cub.2017.10.009. PubMed PMID: 29103937; PubMed Central PMCID: PMCPMC5698132.

6. Liu Y, Sun J. Detection of pathogens and regulation of immunity by the *Caenorhabditis elegans* nervous system. mBio. 2021;12(2). Epub 2021/04/01. doi: 10.1128/mBio.02301-20. PubMed PMID: 33785621; PubMed Central PMCID: PMCPMC8092265.

7. Pukkila-Worley R. Surveillance immunity: an emerging paradigm of innate defense activation in *Caenorhabditis elegans*. PLoS Pathog. 2016;12(9):e1005795. Epub 2016/09/16. doi: 10.1371/journal.ppat.1005795. PubMed PMID: 27631629; PubMed Central PMCID: PMCPMC5025020.

8. Dunbar TL, Yan Z, Balla KM, Smelkinson MG, Troemel ER. *C. elegans* detects pathogen-induced translational inhibition to activate immune signaling. Cell Host Microbe. 2012;11(4):375–86. Epub 2012/04/24. doi: 10.1016/j.chom.2012.02.008. PubMed PMID: 22520465; PubMed Central PMCID: PMCPMC3334869.

9. McEwan DL, Kirienko NV, Ausubel FM. Host translational inhibition by *Pseudomonas aeruginosa* exotoxin A triggers an immune response in *Caenorhabditis elegans*. Cell Host Microbe. 2012;11(4):364–74. Epub 2012/04/24. doi: 10.1016/j.chom.2012.02.007. PubMed PMID: 22520464; PubMed Central PMCID: PMCPMC3334877.

10. Melo JA, Ruvkun G. Inactivation of conserved *C. elegans* genes engages pathogen- and xenobiotic-associated defenses. Cell. 2012;149(2):452–66. Epub 2012/04/17. doi: 10.1016/j.cell.2012.02.050. PubMed PMID: 22500807; PubMed Central PMCID: PMCPMC3613046.

11. Reddy KC, Dunbar TL, Nargund AM, Haynes CM, Troemel ER. The *C. elegans* CCAAT-enhancer-binding protein gamma is required for surveillance immunity. Cell Rep. 2016;14(7):1581–9. Epub 2016/02/16. doi: 10.1016/j.celrep.2016.01.055. PubMed PMID: 26876169; PubMed Central PMCID: PMCPMC4767654.

12. Pellegrino MW, Nargund AM, Kirienko NV, Gillis R, Fiorese CJ, Haynes CM. Mitochondrial UPR-regulated innate immunity provides resistance to pathogen infection. Nature. 2014;516(7531):414-7. Epub 2014/10/03. doi: 10.1038/nature13818. PubMed PMID: 25274306; PubMed Central PMCID: PMCPMC4270954.

13. Tjahjono E, Kirienko NV. A conserved mitochondrial surveillance pathway is required for defense against *Pseudomonas aeruginosa*. PLoS Genet. 2017;13(6):e1006876. Epub 2017/07/01. doi: 10.1371/journal.pgen.1006876. PubMed PMID: 28662060; PubMed Central PMCID: PMCPMC5510899.

14. Tecle E, Chhan CB, Franklin L, Underwood RS, Hanna-Rose W, Troemel ER. The purine nucleoside phosphorylase *pnp-1* regulates epithelial cell resistance to infection in *C. elegans*. PLoS Pathog. 2021;17(4):e1009350. Epub 2021/04/21. doi: 10.1371/journal.ppat.1009350. PubMed PMID: 33878133; PubMed Central PMCID: PMCPMC8087013.

15. Reddy KC, Dror T, Underwood RS, Osman GA, Elder CR, Desjardins CA, et al. Antagonistic paralogs control a switch between growth and pathogen resistance in *C. elegans*. PLoS Pathog. 2019;15(1):e1007528. Epub 2019/01/15. doi: 10.1371/journal.ppat.1007528. PubMed PMID: 30640956; PubMed Central PMCID: PMCPMC6347328.

16. Blackwell TK, Steinbaugh MJ, Hourihan JM, Ewald CY, Isik M. SKN-1/Nrf, stress responses, and aging in *Caenorhabditis elegans*. Free Radic Biol Med. 2015;88(Pt B):290–301. Epub 2015/08/02. doi: 10.1016/j.freeradbiomed.2015.06.008. PubMed PMID: 26232625; PubMed Central PMCID: PMCPMC4809198.

17. Lehrbach NJ, Breen PC, Ruvkun G. Protein sequence editing of SKN-1A/Nrf1 by peptide:N-glycanase controls proteasome gene expression. Cell. 2019;177(3):737–50 e15. Epub 2019/04/20. doi: 10.1016/j.cell.2019.03.035. PubMed PMID: 31002798; PubMed Central PMCID: PMCPMC6574124.

18. Radhakrishnan SK, Lee CS, Young P, Beskow A, Chan JY, Deshaies RJ. Transcription factor Nrf1 mediates the proteasome recovery pathway after proteasome inhibition in mammalian cells. Mol Cell. 2010;38(1):17–28. Epub 2010/04/14. doi: 10.1016/j.molcel.2010.02.029. PubMed PMID: 20385086; PubMed Central PMCID: PMCPMC2874685.

19. Li X, Matilainen O, Jin C, Glover-Cutter KM, Holmberg CI, Blackwell TK. Specific SKN-1/Nrf stress responses to perturbations in translation elongation and proteasome activity. PLoS Genet. 2011;7(6):e1002119. Epub 2011/06/23. doi: 10.1371/journal.pgen.1002119. PubMed PMID: 21695230; PubMed Central PMCID: PMCPMC3111486.

20. An JH, Blackwell TK. SKN-1 links *C. elegans* mesendodermal specification to a conserved oxidative stress response. Genes Dev. 2003;17(15):1882–93. Epub 2003/07/19. doi: 10.1101/gad.1107803. PubMed PMID: 12869585; PubMed Central PMCID: PMCPMC196237.

21. Hoeven R, McCallum KC, Cruz MR, Garsin DA. Ce-Duox1/BLI-3 generated reactive oxygen species trigger protective SKN-1 activity via p38 MAPK signaling during infection in *C. elegans*. PLoS Pathog. 2011;7(12):e1002453. Epub 2012/01/05. doi: 10.1371/journal.ppat.1002453. PubMed PMID: 22216003; PubMed Central PMCID: PMCPMC3245310.

22. Papp D, Csermely P, Soti C. A role for SKN-1/Nrf in pathogen resistance and immunosenescence in *Caenorhabditis elegans*. PLoS Pathog. 2012;8(4):e1002673. Epub 2012/05/12. doi: 10.1371/journal.ppat.1002673. PubMed PMID: 22577361; PubMed Central PMCID: PMCPMC3343120.

23. Wu C, Karakuzu O, Garsin DA. Tribbles pseudokinase NIPI-3 regulates intestinal immunity in *Caenorhabditis elegans* by controlling SKN-1/Nrf activity. Cell Rep. 2021;36(7):109529. doi: 10.1016/j.celrep.2021.109529. PubMed PMID: 34407394; PubMed Central PMCID: PMCPMC8393239.

24. Khush RS, Cornwell WD, Uram JN, Lemaitre B. A ubiquitin-proteasome pathway represses the *Drosophila* immune deficiency signaling cascade. Curr Biol. 2002;12(20):1728–37. Epub 2002/10/29. doi: 10.1016/s0960-9822(02)01214-9. PubMed PMID: 12401167.

25. Engel E, Viargues P, Mortier M, Taillebourg E, Coute Y, Thevenon D, et al. Identifying USPs regulating immune signals in *Drosophila:* USP2 deubiquitinates Imd and promotes its degradation by interacting with the proteasome. Cell Commun Signal. 2014;12:41. Epub 2014/07/17. doi: 10.1186/s12964-014-0041-2. PubMed PMID: 25027767; PubMed Central PMCID: PMCPMC4140012.

26. Ebstein F, Poli Harlowe MC, Studencka-Turski M, Kruger E. Contribution of the unfolded protein response (UPR) to the pathogenesis of proteasome-associated autoinflammatory syndromes (PRAAS). Front Immunol. 2019;10:2756. Epub 2019/12/13. doi: 10.3389/fimmu.2019.02756. PubMed PMID: 31827472; PubMed Central PMCID: PMCPMC6890838.

27. Brehm A, Liu Y, Sheikh A, Marrero B, Omoyinmi E, Zhou Q, et al. Additive loss-of-function proteasome subunit mutations in CANDLE/PRAAS patients promote type I IFN production. J Clin Invest. 2015;125(11):4196–211. Epub 2015/11/03. doi: 10.1172/JCI81260. PubMed PMID: 26524591; PubMed Central PMCID: PMCPMC4639987.

28. Gang SS, Grover M, Reddy KC, Raman D, Chang YT, Ekiert DC, et al. A *pals-25* gain-of-function allele triggers systemic resistance against natural pathogens of *C. elegans*. PLoS Genet. 2022;18(10):e1010314. Epub 2022/10/04. doi: 10.1371/journal.pgen.1010314. PubMed PMID: 36191002; PubMed Central PMCID: PMCPMC9560605.

29. Lehrbach NJ, Ruvkun G. Proteasome dysfunction triggers activation of SKN-1A/Nrf1 by the aspartic protease DDI-1. Elife. 2016;5. Epub 2016/08/17. doi: 10.7554/eLife.17721. PubMed PMID: 27528192; PubMed Central PMCID: PMCPMC4987142.

30. Consortium CeDM. large-scale screening for targeted knockouts in the *Caenorhabditis elegans* genome. G3 (Bethesda). 2012;2(11):1415-25. Epub 2012/11/23. doi: 10.1534/g3.112.003830. PubMed PMID: 23173093; PubMed Central PMCID: PMCPMC3484672.

31. Choe KP, Przybysz AJ, Strange K. The WD40 repeat protein WDR-23 functions with the CUL4/DDB1 ubiquitin ligase to regulate nuclear abundance and activity of SKN-1 in *Caenorhabditis elegans*. Mol Cell Biol. 2009;29(10):2704–15. Epub 2009/03/11. doi: 10.1128/MCB.01811-08. PubMed PMID: 19273594; PubMed Central PMCID: PMCPMC2682033.

32. Leung CK, Hasegawa K, Wang Y, Deonarine A, Tang L, Miwa J, et al. Direct interaction between the WD40 repeat protein WDR-23 and SKN-1/Nrf inhibits binding to target DNA. Mol Cell Biol. 2014;34(16):3156–67. Epub 2014/06/11. doi: 10.1128/MCB.00114-14. PubMed PMID: 24912676; PubMed Central PMCID: PMCPMC4135592.

33. Lehrbach NJ, Ruvkun G. Endoplasmic reticulum-associated SKN-1A/Nrf1 mediates a cytoplasmic unfolded protein response and promotes longevity. Elife. 2019;8. Epub 2019/04/12. doi: 10.7554/eLife.44425. PubMed PMID: 30973820; PubMed Central PMCID: PMCPMC6459674.

34. Olaitan AO, Aballay A. Non-proteolytic activity of 19S proteasome subunit RPT-6 regulates GATA transcription during response to infection. PLoS Genet. 2018;14(9):e1007693. Epub 2018/09/29. doi: 10.1371/journal.pgen.1007693. PubMed PMID: 30265660; PubMed Central PMCID: PMCPMC6179307.

35. Zhang P, Qu H-Y, Wu Z, Na H, Hourihan J, Zhang F, et al. ERK signaling licenses SKN-1A/NRF1 for proteasome production and proteasomal stress resistance. bioRxiv. 2021:2021.01.04.425272. doi: 10.1101/2021.01.04.425272.

36. Calixto A, Chelur D, Topalidou I, Chen X, Chalfie M. Enhanced neuronal RNAi in *C. elegans* using SID-1. Nat Methods. 2010;7(7):554–9. Epub 2010/06/01. doi: 10.1038/nmeth.1463. PubMed PMID: 20512143; PubMed Central PMCID: PMCPMC2894993.

37. Espelt MV, Estevez AY, Yin X, Strange K. Oscillatory Ca2+ signaling in the isolated *Caenorhabditis elegans* intestine: role of the inositol-1,4,5-trisphosphate receptor and phospholipases C beta and gamma. J Gen Physiol. 2005;126(4):379-92. Epub 2005/09/28. doi: 10.1085/jgp.200509355. PubMed PMID: 16186564; PubMed Central PMCID: PMCPMC2266627.

38. Drury F, Grover M, Hintze M, Saunders J, Fasseas MK, Constantinou C, et al. A receptor tyrosine kinase regulated by the transcription factor VAB-3/PAX6 pairs with a pseudokinase to trigger immune signalling upon oomycete recognition in *C. elegans*. bioRxiv. 2022. doi: 10.1101/2022.12.15.520564.

39. Segref A, Kevei E, Pokrzywa W, Schmeisser K, Mansfeld J, Livnat-Levanon N, et al. Pathogenesis of human mitochondrial diseases is modulated by reduced activity of the ubiquitin/proteasome system. Cell Metab. 2014;19(4):642–52. Epub 2014/04/08. doi: 10.1016/j.cmet.2014.01.016. PubMed PMID: 24703696.

40. Anderson RT, Bradley TA, Smith DM. Hyperactivation of the proteasome in *Caenorhabditis elegans* protects against proteotoxic stress and extends lifespan. J Biol Chem. 2022;298(10):102415. Epub 2022/08/26. doi: 10.1016/j.jbc.2022.102415. PubMed PMID: 36007615; PubMed Central PMCID: PMCPMC9486566.

41. Troemel ER, Felix MA, Whiteman NK, Barriere A, Ausubel FM. Microsporidia are natural intracellular parasites of the nematode *Caenorhabditis elegans*. PLoS Biol. 2008;6(12):2736–52. Epub 2008/12/17. doi: 10.1371/journal.pbio.0060309. PubMed PMID: 19071962; PubMed Central PMCID: PMCPMC2596862.

42. Tan MW, Mahajan-Miklos S, Ausubel FM. Killing of *Caenorhabditis elegans* by *Pseudomonas aeruginosa* used to model mammalian bacterial pathogenesis. Proc Natl Acad Sci U S A. 1999;96(2):715–20. Epub 1999/01/20. doi: 10.1073/pnas.96.2.715. PubMed PMID: 9892699; PubMed Central PMCID: PMCPMC15202.

43. Chen DD, Jiang JY, Lu LF, Zhang C, Zhou XY, Li ZC, et al. Zebrafish Uba1 degrades IRF3 through K48-linked ubiquitination to inhibit IFN production. J Immunol. 2021;207(2):512–22. Epub 2021/07/02. doi: 10.4049/jimmunol.2100125. PubMed PMID: 34193603.

44. and AJZ, Harshman LG. The physiology of life history trade-offs in animals. Annual Review of Ecology and Systematics. 2001;32(1):95–126. doi: 10.1146/annurev.ecolsys.32.081501.114006.

45. Lažetić V, Blanchard MJ, Bui T, Troemel ER. Multiple *pals* gene modules control a balance between immunity and development in *Caenorhabditis elegans*. PLOS Pathogens. 2023;19(7):e1011120. doi: 10.1371/journal.ppat.1011120.

46. Yan K, Zhang J, Lee PY, Tao P, Wang J, Wang S, et al. Haploinsufficiency of PSMD12 causes proteasome dysfunction and subclinical autoinflammation. Arthritis Rheumatol. 2022;74(6):1083–90. Epub 2022/01/27. doi: 10.1002/art.42070. PubMed PMID: 35080150; PubMed Central PMCID: PMCPMC9321778.

47. Tataridas-Pallas N, Thompson MA, Howard A, Brown I, Ezcurra M, Wu Z, et al. Neuronal SKN-1B modulates nutritional signalling pathways and mitochondrial networks to control satiety. PLoS Genet. 2021;17(3):e1009358. Epub 2021/03/05. doi: 10.1371/journal.pgen.1009358. PubMed PMID: 33661901; PubMed Central PMCID: PMCPMC7932105.

48. Cetin G, Studencka-Turski M, Venz S, Schormann E, Junker H, Hammer E, et al. Immunoproteasomes control activation of innate immune signaling and microglial function. Front Immunol. 2022;13:982786. Epub 2022/10/25. doi: 10.3389/fimmu.2022.982786. PubMed PMID: 36275769; PubMed Central PMCID: PMCPMC9584546.

49. Waad Sadiq Z, Brioli A, Al-Abdulla R, Cetin G, Schutt J, Murua Escobar H, et al. Immunogenic cell death triggered by impaired deubiquitination in multiple myeloma relies on dysregulated type I interferon signaling. Front Immunol. 2023;14:982720. Epub 2023/03/21. doi: 10.3389/fimmu.2023.982720. PubMed PMID: 36936919; PubMed Central PMCID: PMCPMC10018035.

50. Lazetic V, Wu F, Cohen LB, Reddy KC, Chang YT, Gang SS, et al. The transcription factor ZIP-1 promotes resistance to intracellular infection in *Caenorhabditis elegans*. Nat Commun. 2022;13(1):17. Epub 2022/01/12. doi: 10.1038/s41467-021-27621-w. PubMed PMID: 35013162; PubMed Central PMCID: PMCPMC8748929.

51. Stiernagle T. Maintenance of *C. elegans*. WormBook. 2006:1–11. Epub 20060211. doi: 10.1895/wormbook.1.101.1. PubMed PMID: 18050451; PubMed Central PMCID: PMCPMC4781397.

52. Minevich G, Park DS, Blankenberg D, Poole RJ, Hobert O. CloudMap: a cloud-based pipeline for analysis of mutant genome sequences. Genetics. 2012;192(4):1249–69. Epub 2012/10/12. doi: 10.1534/genetics.112.144204. PubMed PMID: 23051646; PubMed Central PMCID: PMCPMC3512137.

53. Estes KA, Szumowski SC, Troemel ER. Non-lytic, actin-based exit of intracellular parasites from *C. elegans* intestinal cells. PLoS Pathog. 2011;7(9):e1002227. Epub 2011/09/29. doi: 10.1371/journal.ppat.1002227. PubMed PMID: 21949650; PubMed Central PMCID: PMCPMC3174248.

54. Bray NL, Pimentel H, Melsted P, Pachter L. Near-optimal probabilistic RNA-seq quantification. Nat Biotechnol. 2016;34(5):525–7. Epub 20160404. doi: 10.1038/nbt.3519. PubMed PMID: 27043002.

55. Pimentel H, Bray NL, Puente S, Melsted P, Pachter L. Differential analysis of RNA-seq incorporating quantification uncertainty. Nat Methods. 2017;14(7):687–90. Epub 20170605. doi: 10.1038/nmeth.4324. PubMed PMID: 28581496.

56. Timmons L, Court DL, Fire A. Ingestion of bacterially expressed dsRNAs can produce specific and potent genetic interference in *Caenorhabditis elegans*. Gene. 2001;263(1-2):103–12. Epub 2001/02/27. doi: 10.1016/s0378-1119(00)00579-5. PubMed PMID: 11223248.

57. Reboul J, Vaglio P, Tzellas N, Thierry-Mieg N, Moore T, Jackson C, et al. Open-reading-frame sequence tags (OSTs) support the existence of at least 17,300 genes in C. elegans. Nat Genet. 2001;27(3):332–6. Epub 2001/03/10. doi: 10.1038/85913. PubMed PMID: 11242119.

58. Kamath RS, Fraser AG, Dong Y, Poulin G, Durbin R, Gotta M, et al. Systematic functional analysis of the *Caenorhabditis elegans* genome using RNAi. Nature. 2003;421(6920):231-7. Epub 2003/01/17. doi: 10.1038/nature01278. PubMed PMID: 12529635.

59. Hintze M, Koneru SL, Gilbert SPR, Katsanos D, Lambert J, Barkoulas M. A cell fate switch in the *Caenorhabditis elegans* seam cell lineage occurs through modulation of the Wnt asymmetry pathway in response to temperature increase. Genetics. 2020;214(4):927–39. Epub 20200127. doi: 10.1534/genetics.119.302896. PubMed PMID: 31988193; PubMed Central PMCID: PMCPMC7153939.

